# Sequential deregulation of histone marks, chromatin accessibility and gene expression in response to PROTAC-induced degradation of ASH2L

**DOI:** 10.1101/2023.09.08.556832

**Authors:** Mirna Barsoum, Roksaneh Sayadi-Boroujeni, Alexander T. Stenzel, Philip Bussmann, Juliane Lüscher-Firzlaff, Bernhard Lüscher

**Author notes:** Contributed equally (Mirna Barsoum, Roksaneh Sayadi-Boroujeni). Correspondence to M.B. and B.L.

## Abstract

The trithorax protein ASH2L is essential for organismal and tissue development. As a subunit of COMPASS/KMT2 complexes, ASH2L is necessary for methylation of histone H3 lysine 4 (H3K4). Mono- and trimethylation at this site mark active enhancers and promoters, respectively, although the molecular relevance of H3K4 methylation is only partially understood. Due to the importance of ASH2L in all 6 COMPASS-like complexes and its long half-life, it has been difficult to define direct consequences. To overcome this limitation, we employed a PROTAC system, which allows the rapid degradation of ASH2L and addressing direct effects. Loss of ASH2L resulted in rapid inhibition of proliferation of mouse embryo fibroblasts. Shortly after ASH2L degradation, H3K4me3 decreased with its half-life revealing considerable variability between promoters. Subsequently, H3K4me1 increased at promoters and decreased at some enhancers. H3K27ac and H3K27me3, histone marks closely linked to H3K4 methylation, were affected with considerable delay. In parallel, chromatin compaction increased at promoters. Of note, nascent gene transcription was not affected early but overall RNA expression was deregulated late after ASH2L loss. Together, these findings suggest that downstream effects are ordered but relatively slow, despite the rapid loss of ASH2L and inactivation of KMT2 complexes. It appears that the systems that control gene transcription are well buffered and strong effects are only beginning to unfold after considerable delay.

## Introduction

In eukaryotic cells, the genome is organized as chromatin to control the access to and the use of the DNA in processes such as transcription, replication and repair. The smallest unit of chromatin is the nucleosome, which is composed of four core histones, H2A, H2B, H3 and H4 or variants thereof, and 147 base pairs of DNA^1^. Linker DNA of various length connects individual nucleosomes. Higher order chromatin organization involves loops, topologically associated domains or TADs, and A/B compartments, defined at least in part by factors such as CTCF and cohesin^2,3^. The core of nucleosomes is formed by the globular regions of histones, while the N-terminal tails protrude, which makes them accessible to a wide range of post-translational modifications (PTMs)^4,5^. These are the product of a large panel of enzymes that include writers such as methyltransferases and acetyltransferases and the corresponding erasers, thereby controlling access to nucleosomes, DNA and more general chromatin. Sequence-specific transcription factors are key to direct and assemble these enzymes to distinct regions in chromatin and to coordinate the regulation of gene transcription^6,7^.

Reversible methylation of histone H3 at lysine 4 (H3K4) has been linked to gene transcription^8–11^. Lysines can be mono-, di- or tri-methylated (Kme1-3), thereby altering the spectrum of reader molecules that are able to interact. While H3K4me1 is primarily located at enhancers, H3K4me3 is a mark of open chromatin at promoters. Together with H3K27 acetylation (H3K27ac), these marks define active enhancers and promoters, respectively^12–15^. In contrast, H3K4me3 in combination with H3K27me3 establish bivalent chromatin, which is linked to poised promoters^16,17^. Multiple readers have been described, which are thought to convey information encoded in H3K4 methylation, whose effects include chromatin remodeling, RNA polymerase (RNAPII) loading, and H3K4 methylation amplification^18,19^.

H3K4 methylation is catalyzed by COMPASS (complex of proteins associated with Set1)^9–11^. This complex, originally defined in yeast^20^, exists in 6 versions in mammals, defined by 6 different catalytic subunits, KMT2A-D, F and G (MLL1-4, SET1A and B, respectively). All 6 versions contain a core complex of 4 proteins, WDR5, RBBP5, ASH2L and 2 copies of DPY30, the so-called WRAD complex, which is necessary for efficient methyltransferase activity^21–26^. Additional subunits have been described that are specific for certain KMT2 complexes^10,27^. As far as studied, all the subunits of KMT2 complexes are essential for proper cell functioning, particularly the WRAD subunits, as their deletion results in distinct developmental defects in model organisms. This poses challenges for WRAD subunit analyses as broad effects on chromatin and gene expression are expected and observed, resulting in complex phenotypes. Also, many of the subunits are mutated or their expression deregulated in diseases, including cancer, neurodegeneration and complex syndromes^27–29^. Of note is that the WRAD complex interacts with multiple sequence-specific transcription factors and thus appears to be important for recruiting COMPASS-like complexes to specific sites in chromatin^27^. In addition, these complexes interact with the RNAPII complex and with CpG islands^11,30^. Thus, the different KMT2 complexes are assembled from common and selective subunits that are important for optimal catalytic activity and chromatin localization.

ASH2L is necessary for organismal development^31,32^. Moreover, we have previously observed that deletion of *Ash2l* in the hematopoietic system prevents proliferation and differentiation of hematopoietic cells, ultimately resulting in the death of the animals^33^. Of note is that the loss of Dpy30 results in a very similar phenotype^34,35^, suggesting that the main functions of Ash2l and Dpy30 are associated with the WRAD complex and thus with KMT2 complexes. We identified ASH2L as an interaction partner of the oncoprotein MYC^36,37^, indicating that this transcription factor can recruit KMT2 complexes. Indeed, sequential chromatin immunoprecipitation (ChIP) experiments documented that the two proteins can co-localize to known MYC response elements and that binding of MYC is associated with increased H3K4me3^37^. Mechanistically, downregulation or loss of ASH2L provokes a decrease of H3K4me3 at promoters, associated with altered gene transcription^33,37–40^. Somewhat counterintuitive, the loss of Ash2l and the decrease in H3K4me3 at promoters, both linked positively to gene expression, cause both repression and activation of gene transcription. We have argued previously that activation might well be a secondary effect, for example when a transcriptional repressor is no longer expressed^39,40^. This might be particularly relevant in experimental settings that are characterized by a slow response such as upon applying siRNA or using classical recombination (see also below). Together, these studies suggested that in cells ASH2L is necessary for efficient H3K4me3, affecting gene expression, findings that are consistent with a determining function of H3K4me3 for promoter activity.

Our previous work, based on efficient and fast recombination of the floxed *Ash2l* alleles, was hampered due to the long half-life of ASH2L proteins, associated with a slow development of phenotypes. For example, in mouse embryo fibroblasts (MEFs) substantial effects on H3K4 methylation and gene expression were seen only after several days upon loss of Ash2l^39,40^. Similarly, the manifestation on cell proliferation inhibition, cell cycle arrest and induction of senescence occurred after 5 days or later. Thus, due to the sluggishness of the system, it has been difficult to separate and distinguish primary from secondary and even tertiary effects. Therefore, we developed a system, in which we were able to deregulate ASH2L rapidly by expressing an FKBP-ASH2L fusion protein that is sensitive to a proteolysis targeting chimera (PROTAC). We expressed FKBP-ASH2L in MEF cells with floxed *Ash2l* alleles, deleted the endogenous alleles, rendering the cells dependent on the introduced fusion protein. The PROTAC dTAG-13 binds to FKBP and Cereblon, a subunit of an E3 ligase, and destines FKBP fusion proteins for degradation ^41^. The rapid loss of FKBP-ASH2L inhibits cell proliferation and promotes a consecutive modulation of histone marks at both H3K4 and H3K27, alters the accessibility of chromatin, and deregulates gene expression.

## Results

### Loss of ASH2L prevents cell proliferation

We have studied the molecular and cellular consequences of Ash2l loss in mouse embryo fibroblasts (MEFs) with floxed *Ash2l* alleles and an inducible Cre-ER recombinase (iMEF- *Ash2l^fl/fl^*-Cre-ER). While the knockout of *Ash2l* was rapid, the downstream effects, including the decrease in promoter-associated H3K4me3, altered gene expression, and cell cycle and proliferation arrest, were slow likely due to the long half-life of Ash2l^36,39,40^. Thus, to define direct consequences of the loss of Ash2l and to distinguish these from secondary and further downstream effects has been difficult. To overcome this, we established and utilized a PROTAC system (summarized schematically in Fig. 1a). We generated a plasmid that expresses FKBP-F36V fused through a linker with 2 HA-tags to human ASH2L. FKBP-F36V is a mutant version of the prolyl isomerase FKBP12, which has been engineered to accommodate a ligand that cannot bind to the wild-type protein^41–43^. FKBP-HA2-ASH2L can be tied to Cereblon (CRBN), a component of an E3 ubiquitin ligase complex, using the heterobifunctional compound dTAG-13^41^. The construct expressing FKBP-HA2-ASH2L was introduced into the iMEF-*Ash2l^fl/fl^*-Cre-ER cells. Then, exon 4 of the endogenous *Ash2l* was deleted upon activation of Cre-ER and the corresponding RNA could no longer be detected (Supplementary Fig. S1a)^40^.

**Figure 1.**
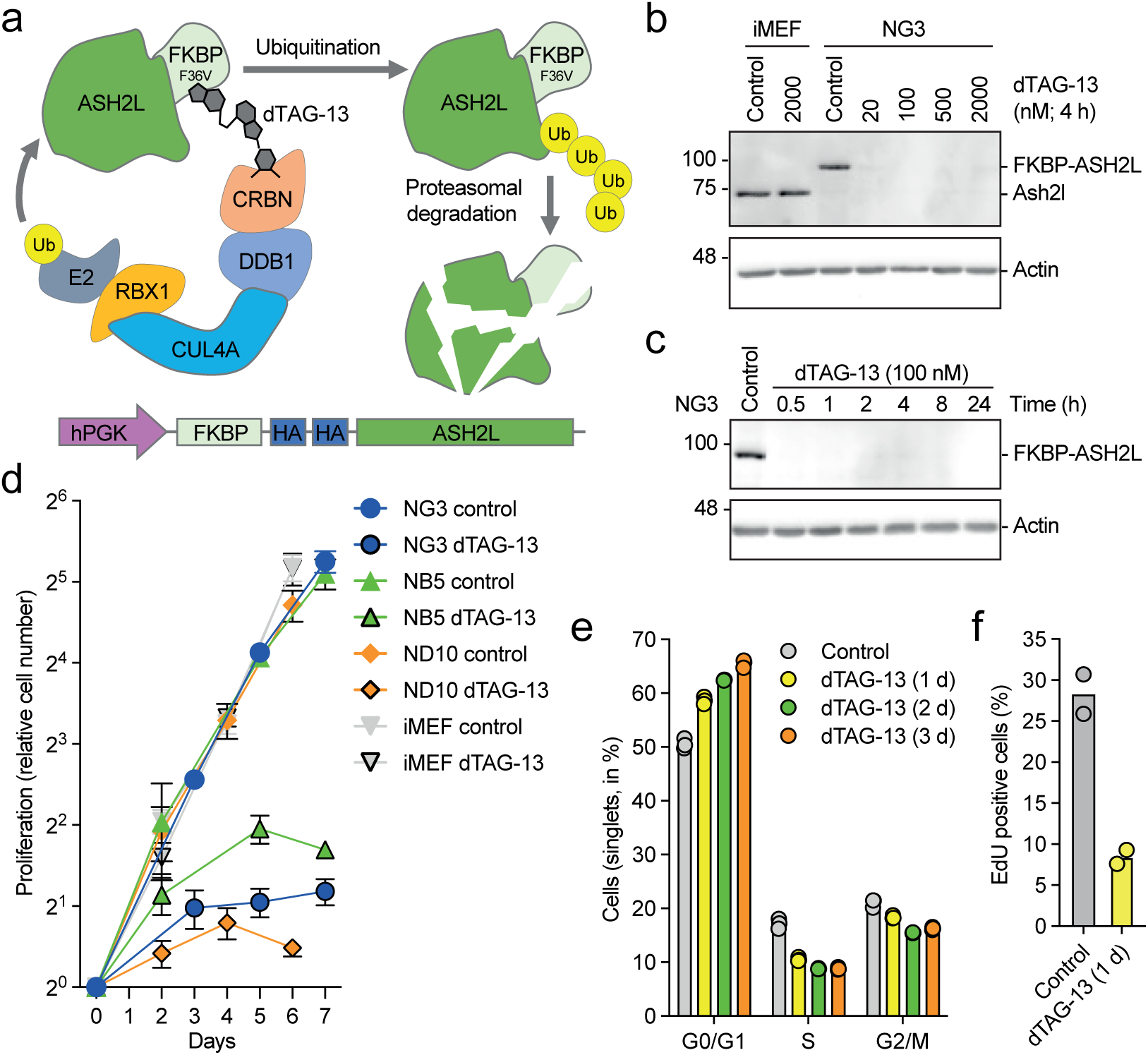
PROTAC-induced rapid degradation of ASH2L inhibits cell proliferation. (a) Schematic summary of dTAG-13 mediated degradation of FKBP-HA2-ASH2L fusion proteins. (b) iMEF control cells and NG3 cells were treated with dTAG-13 as indicated. Total cell lysates were analyzed for Ash2l and FKBP-HA2-ASH2L expression using ASH2L/Ash2l selective antibodies. Actin staining was used as loading control. (c) As in panel B with dTAG-13 treatment for the indicated times. (d) Clones expressing FKBP-HA2-ASH2L fusion proteins (NG3, NB5, ND10) and control cells (iMEF) were treated with or without dTAG13 from day 0. Cells were counted at the indicated time points. Measurement were in triplicates with three biological replicates. Indicated are relative mean values ±SEM. (e) NB3 cells were treated with dTAG-13 for the indicated times. The cells were fixed and stained with Hoechst 33258 and analyzed by flow cytometry. Mean values of two measurements in triplicates are displayed. (f) NB3 cells were treated with or without dTAG-13 for 24 hours. During the final two hours, the cells were incubated with EdU, fixed and stained with AF488 azide. Mean values of two flow cytometry measurements in triplicates are shown.

The expression of the endogenous Ash2l and the FKBP-HA2-ASH2L fusion protein was analyzed in response to dTAG-13 (Fig. 1a). We established three clones, NG3, ND10 and NB5, in which the endogenous Ash2l could no longer be detected (Fig. 1b and Supplementary Fig. S1b and c). The expression of the introduced FKBP-HA2-ASH2L was comparable to the endogenous Ash2l (Fig. 1b). In all three clones the FKBP-HA2-ASH2L protein was sensitive to dTAG-13 and the protein was degraded efficiently by 30 min (Fig. 1b and c, Supplementary Fig. S1b and c). Titration experiments of control lysates suggested that less than 1% of FKBP-HA2-ASH2L remained after 1 hour of dTAG-13 treatment (Supplementary Fig. S1d). Further studies were predominantly performed with NG3 cells.

The loss of Ash2l results in inhibition of proliferation and a block of cell cycle progression in both MEF cells and in hematopoietic multi-potent progenitors^33,40^. In the latter, human ASH2L rescues cell proliferation in tissue culture^33^. Similarly, the FKBP-HA2-ASH2L clones, in the absence of endogenous Ash2l, proliferated comparably to control iMEF cells (Fig. 1d). These cells were unaffected by dTAG-13 (Fig. 1d). In contrast, all three FKBP-HA2-ASH2L clones stopped proliferating rapidly, for example, clone NG3 cells doubled once initially and then cell numbers remained constant (Fig. 1d). A small increase in G0/G1 cells was noted over 3 days, but no distinct cell cycle arrest was observed (Fig. 1e), yet the incorporation of the thymidine analog 5-ethynyl-2’-deoxyuridine (EdU) decreased strongly within 24 hours (Fig. 1f). Thus, no obvious specific checkpoint seemed to be activated but rather supports the notion that the loss of ASH2L generates dependencies on various factors associated with various processes that result in the arrest of cells throughout the cell cycle, similar to what we had observed in MEFs cells upon *Ash2l* knockout^40^. We did not observe any sign of apoptosis or senescence in the first 24 hours. Also, neither p53 expression nor phosphorylation of H2A.X (γ-H2A.X) were induced as might occur upon altered chromatin organization or replication stalling and associated DNA damage (Supplementary Fig. S1e and f). For control, cells were treated with etoposide, which promoted p53 accumulation and H2A.X phosphorylation. Together, these findings reiterate that ASH2L is necessary for cell proliferation.

### ASH2L loss deregulates gene expression

ASH2L is necessary for assembling the WRAD complex, which is required for efficient H3K4 methyltransferase activity of all six KMT2 enzymes and thus for global H3K4 methylation. Indeed, the loss of Ash2l in the iMEF system reduced H3K4 methylation, a process that took several days^39,40^. In the PROTAC system, a rapid decrease of H3K4me3 was measured (Fig. 2a and Supplementary Fig. S1b and c). Assuming that very little KMT2 catalytic activity remains, the decrease in H3K4me3 should depend on demethylases that have been described to remove methylation from H3K4, including KDM5 family members and LSD1^8,44^. Quantification of Western blots and comparing the H3K4me3 signals to total H3 revealed that until 1 hour no decrease in methylation was measured. Subsequently, the decrease was rapid between 1 and 8 hours and considerably slower until 48 hours (Fig. 2b). At this time point, roughly 10% of H3K4me3 was remaining. The overall decrease in H3K4me1 was slower (Fig. 2a), possibly in part due to an increase in mono-methylation at promoters, as discussed below. Further, the sensitivity of H3K4me3 and H3K4me1 to the different demethylases may be distinct^8,44^. The decrease in H3K27ac was also slow, while H3K27me3 did not change (Fig. 2a). Thus, although FKBP-HA2-ASH2L was rapidly lost upon dTAG-13 treatment, the changes of the measured histone marks were considerably delayed.

**Figure 2.**
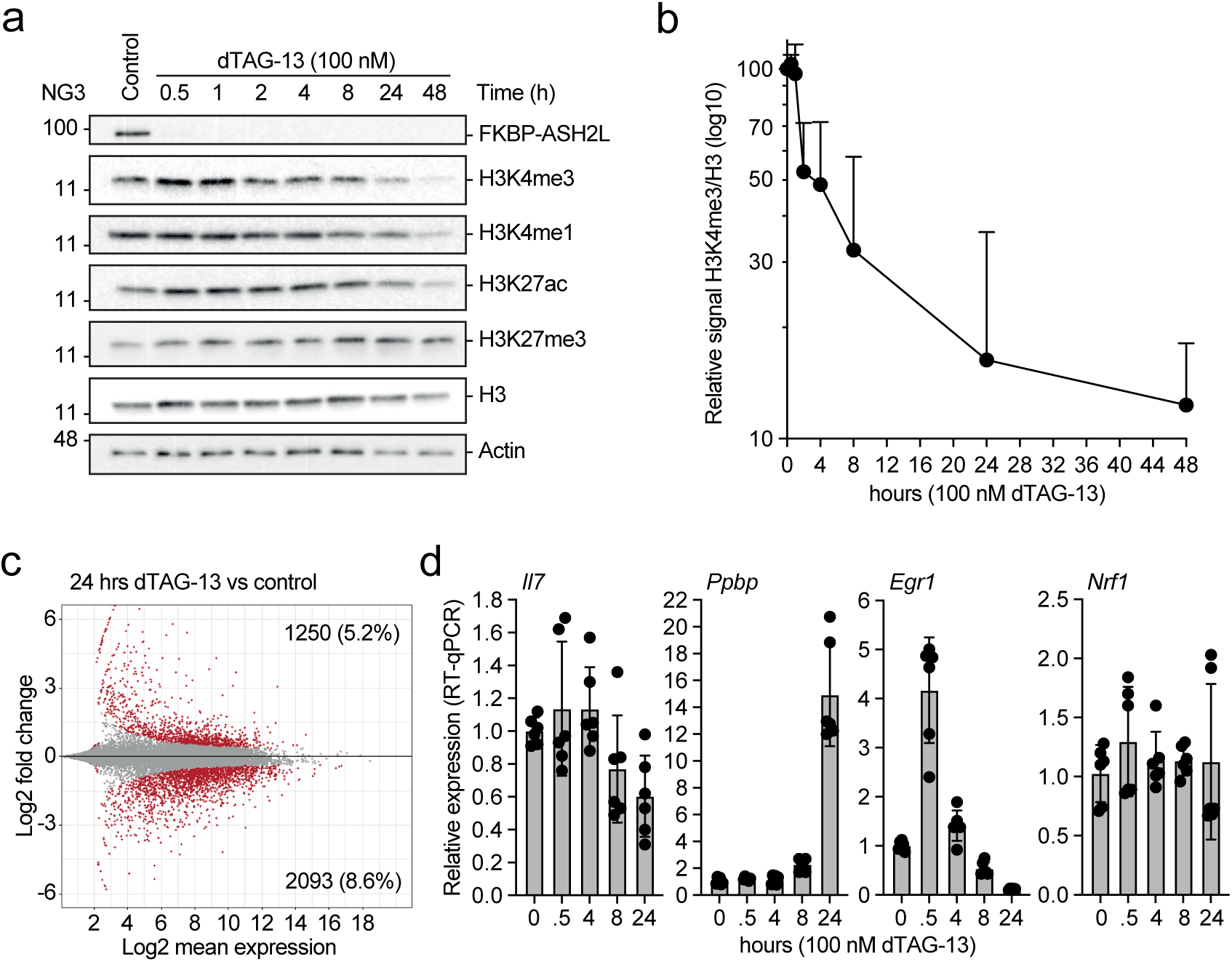
Decrease of H3K4me3 and altered gene expression upon loss of ASH2L. (a) NG3 cells were treated with dTAG-13 for the indicated times. Whole cell lysates were analyzed and the indicated antigens stained on Western blots using selective antibodies. (b) Quantification of H3K4me3 signals compared to total histone H3. Mean of three measurements with SD. (c) NG3 cells were treated with or without dTAG-13 for 24 hours. Whole RNA was extracted and analyzed using 3’mRNA-seq. Displayed is an MA plot. The number of genes that are significantly up- or down-regulated are indicated (red dots: q<0.05). The data summarize two replicates normalized using ERCC spike-in RNA. (d) RT-qPCR analysis of gene expression in response to dTAG-13 treatment. Mean values and standard deviations of 5-6 measurements are displayed.

The decrease of H3K4me3, a histone mark that is predominantly found at active promoters, is expected to alter gene transcription. Upon loss of FKBP-HA2-ASH2L, more than 3000 genes were deregulated by 24 hrs. The majority of genes showed reduced expression, but a substantial number were upregulated (Fig. 2c, Supplementary Table S1). These numbers were higher than what we had observed in the iMEF cells 5 days after *Ash2l* KO^40^. The changes measured were confirmed by RT-qPCR analyses of transcripts that were repressed, induced or remained unchanged (Fig. 2d). Of note is *Egr1*, a gene known to be induced upon various signaling processes, including growth factors, cytokines and different forms of stress^45,46^, which was initially stimulated in response to dTAG-13 treatment and degradation of FKBP-HA2-ASH2L. This suggested that inactivating KMT2 complexes resulted in rapid altered signaling and/or altered sensitivity to signals that control *Egr1* expression.

As expected from the strong inhibition of proliferation, many genes expressing cell cycle relevant proteins, including G1, G1-S, S and G2-M cyclins and replication factors, were downregulated (Supplementary Fig. S1g, Supplementary Table S1). The expression of genes encoding the different Kmt2 methyltransferases, Wdr5, Rbbp5 and Dpy30 was not affected (Supplementary Table S1). Consistent with this finding, the expression of Rbbp5 and Wdr5 was not altered, even after prolonged dTAG-13 treatment (Supplementary Fig. S1h). Thus, it appears that targeting ASH2L does not result in the degradation of other WRAD complex components as might be expected from cross-poly-ubiquitination of the other subunits. It is possible that this occurs but once the existing complexes are depleted, FKBP-HA2-ASH2L might be targeted for degradation before efficient assembly into WRAD complexes can occur.

### Sequential changes in histone marks upon ASH2L loss

To further evaluate the consequences of FKBP-HA2-ASH2L loss, we performed ChIP-seq experiments. Essentially all FKBP-HA2-ASH2L binding sites (6444 sites identified across all samples, Supplementary Table S2a), evaluated using HA-selective antibodies, showed considerably less signal within 1 hour of dTAG-13 treatment (Supplementary Table S2a and b, see also below). For example, FKBP-HA2-ASH2L could no longer be detected at the *Rspo2* and *Zfp503* promoters upon dTAG-13 treatment (Supplementary Fig. S2a). The H3K4 and H3K27 histone marks were measured in time course experiments upon loss of FKBP-HA2-ASH2L (Fig. 3, Supplementary Tables S3-7). The ChIP-seq experiments were performed in duplicates, correlation matrix heat maps demonstrated similarity (Supplementary Fig. S2b and c). We detected H3K4me3 signals at 25431 sites, most of these near transcription start sites (TSS, Supplementary Table S3a). The majority of H3K4me3 binding sites lost signal by 8 hours with little further loss by 16 hours (Fig. 3a, Supplementary Table S3b). We rarely detected an increase in signal. Noticeable was that the downregulated sites were promoter-associated with a strong preference for a small window of ±1 kb around TSSs (Fig. 3b). Despite the overall decrease in H3K4me1 signals, a substantial number of sites showed increased signals (Fig. 3c, Supplementary Table S4). These were almost exclusively located at promoters, while sites with decreased signals were preferentially associated with non-promoter sites (Fig. 3d). This suggested that during the loss of H3K4me3, the H3K4me1 signals increased at least transiently at almost half of the promoters. This increase might contribute to gene repression as suggested earlier^47,48^.

**Figure 3.**
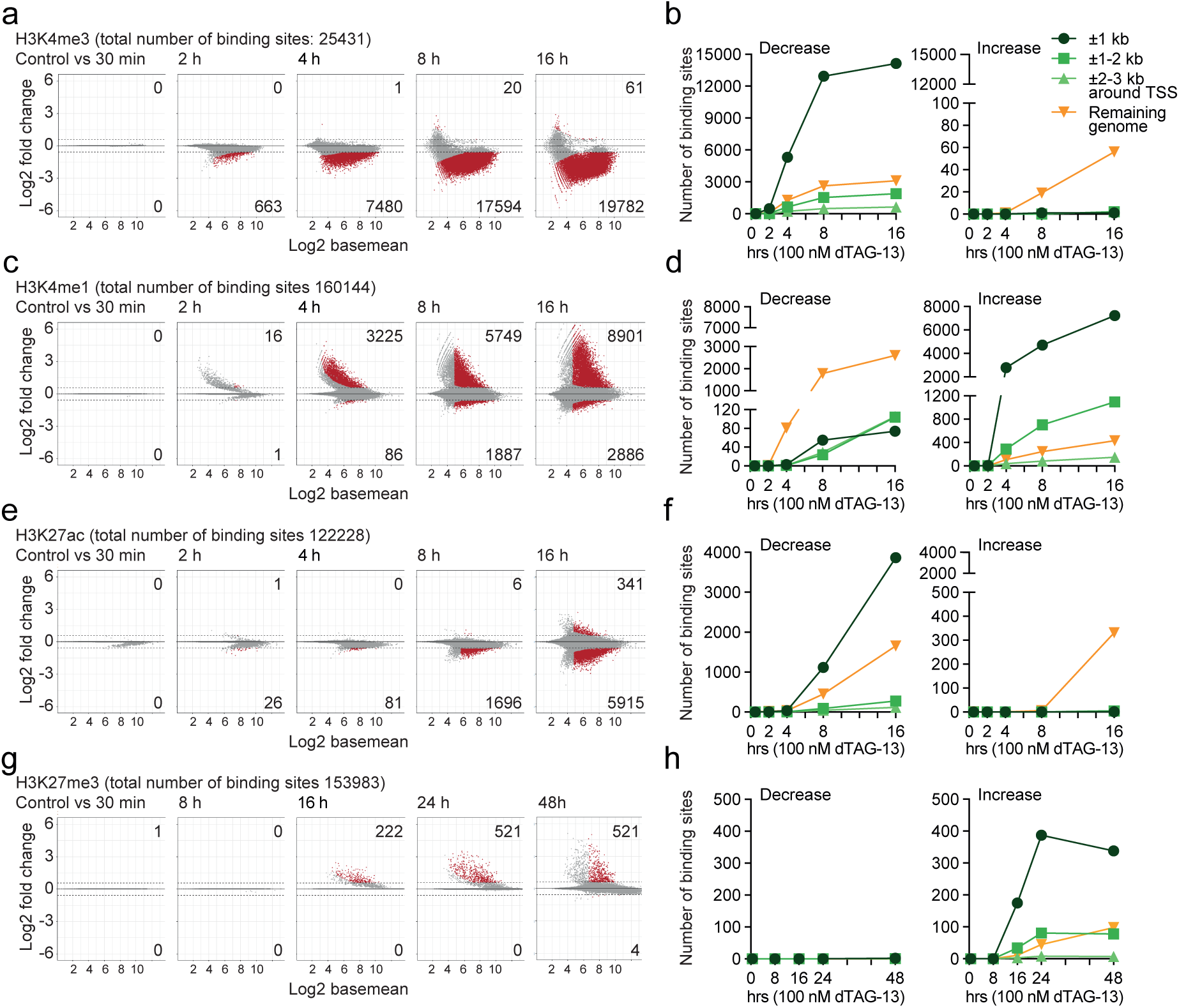
Promoter-associated decrease of H3K4me3 and increase of H3K4me1 upon loss of ASH2L. ChIP-seq experiments of the histone marks H3K4me3 (panels a and b), H3K4me1 (panels c and d), H3K27ac (panels e and f), and H3K27me3 (panels g and h) were performed on cells treated with dTAG-13 for the times indicated (2 biological replicates are summarized; NG3 cells). Panels a, c, e, and g display MA plots. Given are the total number of binding sites identified across all samples and the significantly increased and decreased sites (red dots: q<0.05 and logFC>0.58) upon loss of FKBP-HA2-ASH2L. Panels b, d, f, and h are summary graphs that detail the annotation of significantly changed sites to regions associated with transcriptional start sites (TSS) compared to the remaining genome.

Methylation at H3K4 is connected to acetylation and methylation at H3K27^11,16,49^. Both global H3K27ac and H3K27me3 showed little difference over the first 24 hours upon FKBP-HA2-ASH2L loss (Fig. 2a). H3K27ac decreased slightly by 48 hrs. The ChIP-seq analysis revealed a decrease of H3K27ac signals at roughly 5% of all sites identified after 16 hours (Fig. 3e, Supplementary Fig. S2c, Supplementary Table S5). The majority of these were at promoters and more specifically within ±1 kb of TSSs (Fig. 3f). Sites with increased signals were rare. The findings were verified using ChIP-qPCR. At two selected promoters a decrease of H3K4me3 and H3K27ac was observed, while a tendency to increased H3K4me1 was seen (Supplementary Fig. S2d). Demethylation of H3K4me3 and deacetylation of H3K27ac, particularly at CGI promoters, have been suggested to correlate with an increase of H3K27me3 (Ref^11,16,49^). However, we observed very little effects on H3K27me3 signals (Fig. 3g). At the first time points, no changes in H3K27me3 were detected and only after 16 hrs and beyond a few sites showed enhanced signals (Fig. 3g, Supplementary Table S6 and 7). Despite the small numbers, the majority of changes observed occurred at promoters (Fig. 3h). These effects are also documented with integrative genomics viewer (IGV) browser snapshots comparing the different histone marks at the *Egr1* promoter (Supplementary Fig. S2e). The decrease in H3K4me3 in the promoter region was accompanied by an increase in H3K4me1, a decrease in H3K27ac and an increase in H3K27me3 at late time points. Despite the increase in mRNA after 30 min of dTAG-13 treatment, no obvious changes in histone modifications were apparent at this early time point (Fig. 2d and Supplementary Fig. S2e). The signals for ASH2L were weak and decreased within 1 h of dTAG-13 treatment (Supplementary Fig. S2e). Together, these findings support the conclusions of ordered changes in the studied histone marks as indicated by the genome-wide analyses.

Although the effects on H3K27ac were small, the ChIP-qPCR analysis of selected genes suggested that the decrease in this histone mark can be rapid (Supplementary Fig. S2d). Therefore, we compared the histone marks of promoters that lost H3K4me3 signals either fast or slow (499 and 788, respectively; Fig. 4). H3K4me1 and H3K27ac increased and decreased more rapidly, respectively, in the fast class. While the signals for H3K4me1 picked up at the later time points in the slow class, this was not the case for H3K27ac. For H3K27me3 an overall trend to increased signals at late time points was observed, particularly in the fast class, however, only few of these changes were statistically significant (compare to Fig 3g and h). Furthermore, we divided the promoters with H3K4me3 signals into three equal groups, high, medium and low, according to signal strength (Supplementary Fig. S3), as applied before^39^. We noticed that the decrease over time was comparable. H3K4me1 and H3K27ac increased and decreased, respectively, faster in the high group. Interestingly, the trend to increased H3K27me3 was most prominent in the H3K4me3 high group. Thus, these comparisons suggest that different groups of promoters can be characterized by their varying degrees of H3K4me3 stability and distinct dynamics of other histone marks.

**Figure 4.**
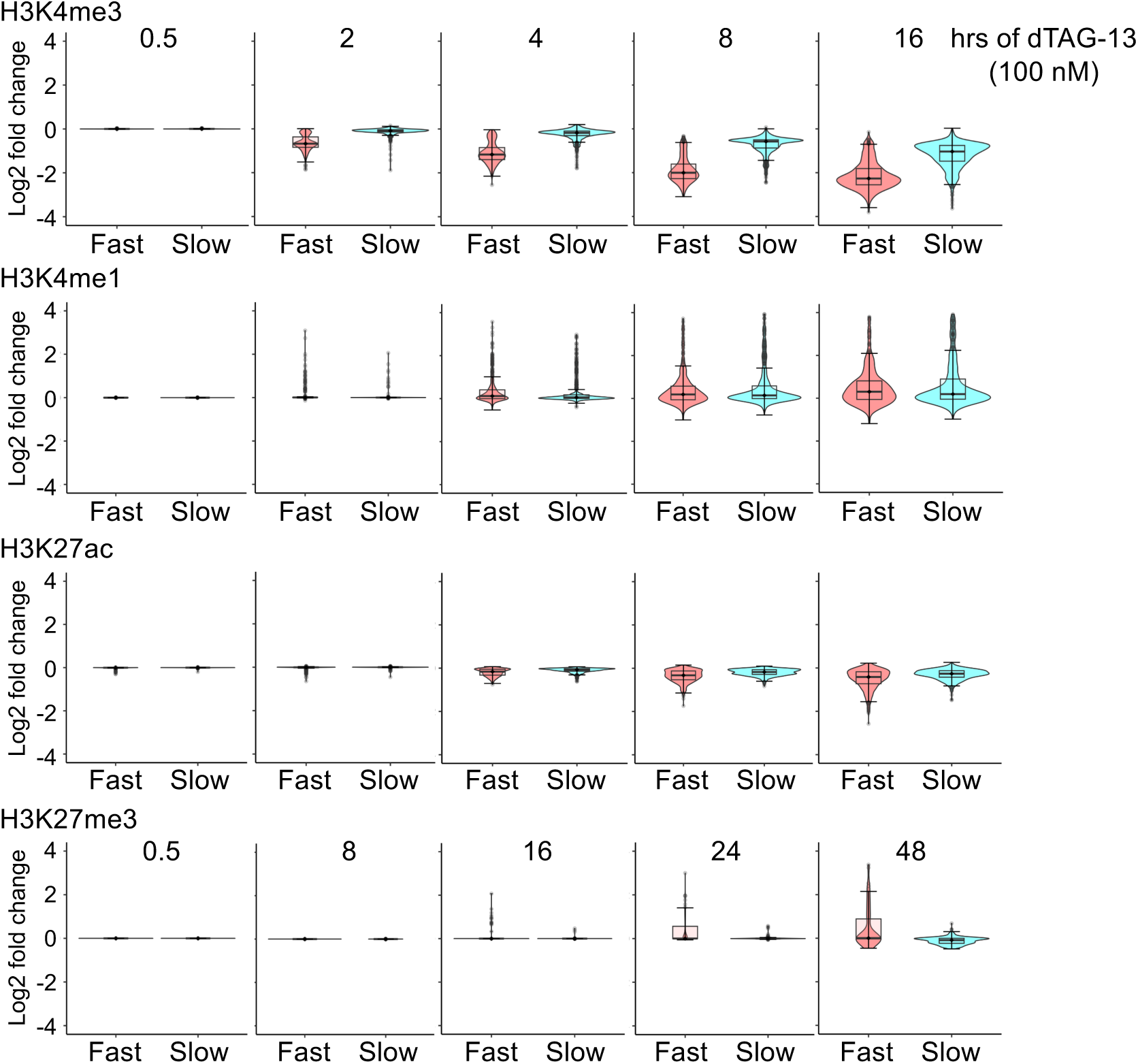
Promoters with a fast loss of H3K4me3 show more rapid alterations of other histone marks. Two classes of promoters were chosen. Promoters that showed significant decrease in H3K4me3 signals either fast (499 at 2 h) or slow (788 at 16 h, excluding promoters that lost H3K4me3 signals after 30 min, 2 h, 4 h or 8 h) (q<0.05; logFC>0.58). A window of ±1 kb around transcriptional start sites was considered. The log2 fold changes of signals of the indicated histone marks at the promoters of these two classes are displayed.

Together, these observations suggest an ordered sequence of events. The rapid loss of FKBP-HA2-ASH2L promotes first a decrease of H3K4me3 at promoters, followed by an increase in H3K4me1 and a decrease in H3K27ac, while changes in H3K27me3 appear late after FKBP-HA2-ASH2L depletion.

### Changes in H3K4 and H3K27 marks occur with a slight preference at CpG island promoters

Because the largest effects were observed at a ±1 kb window centered around TSSs (Fig. 3), we compared all transcripts within the murine genome assembly mm10 with all four histone marks, i.e., H3K4me3, H3K4me1, H3K27ac and H3K27me3, and with FKBP-HA2-ASH2L signals. The ASH2L signals were centered close to TSSs that are positive for H3K4me3 (Fig. 5a and b). The biphasic pattern of H3K4me3 around the TSS with typically more signals 3’ of TSSs was not reflected in the FKBP-HA2-ASH2L binding pattern. Its binding was just slightly 3’ of the TSS (Fig. 5a and b). FKBP-HA2-ASH2L signals were strongly reduced within one hour of dTAG-13 treatment (Fig. 5a and b), consistent with the rapid decrease of the protein measured on Western blots (Fig. 1 and Supplementary Fig. S1). The signals for H3K4me3 and H3K27ac decreased, while H3K4me1 increased at promoters that are labeled with FKBP-HA2-ASH2L in control cells (Fig. 5a-e). The signals for H3K27me3 were very weak and, at the resolution used, no changes could be observed during the first 8 hrs. The number of signals remained low at later time points (Fig. 5a and f).

**Figure 5.**
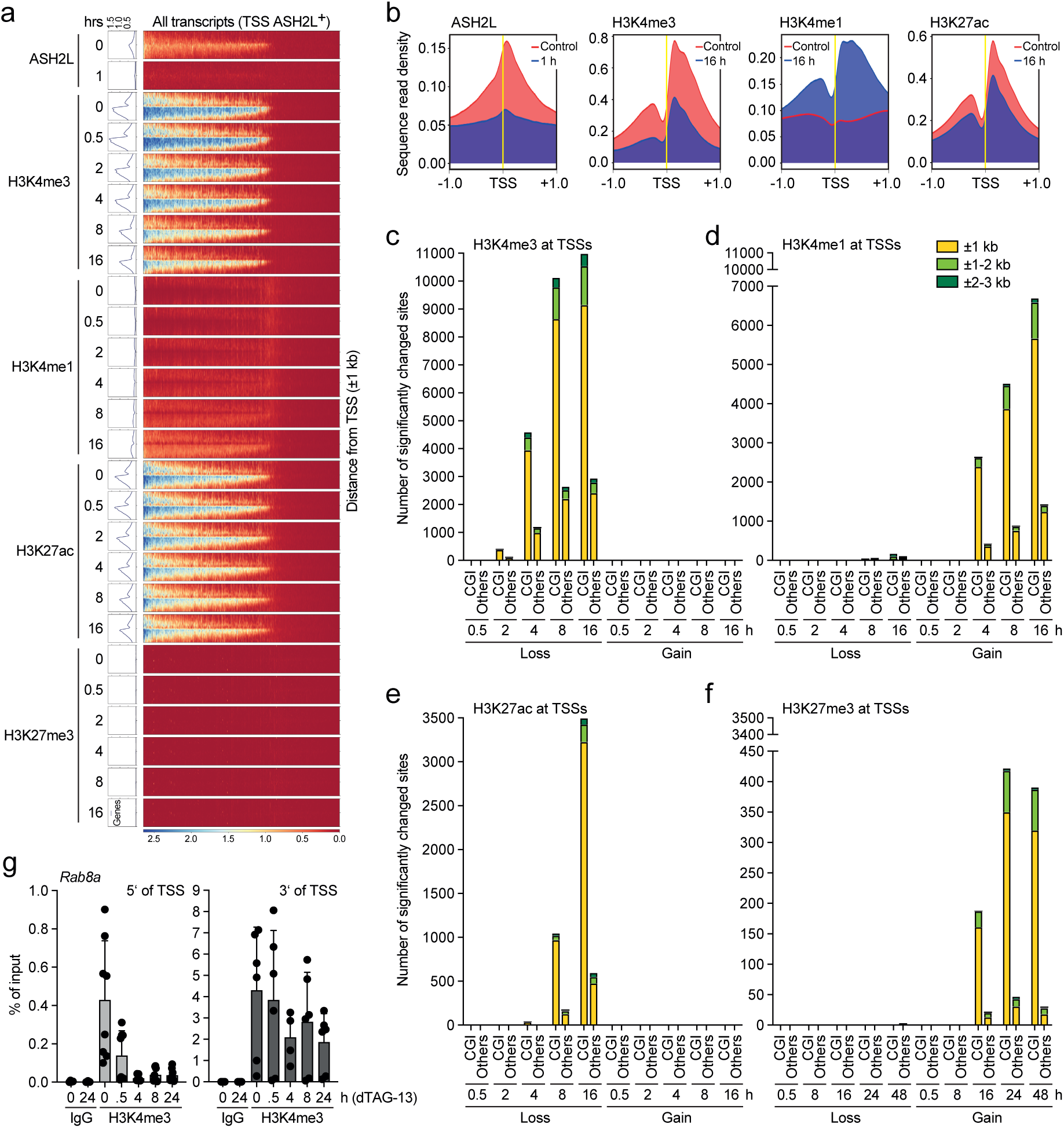
Decrease of H3K4me3 signals upon ASH2L loss is not prevalent at CpG island promoters. (a) Heatmaps and plot profiles generated using DeepTools showing the normalized ChIP-seq signals of the indicated histone marks in response to dTAG-13 treatment at ±1 kb of TSSs of all annotated transcripts in mm10 (normalized using counts per Million). (b) Plot profiles generated using DeepTools show the average normalized signals of the indicated ChIP-seq data as in panel a. (c-f) Lost and gained signals of H3K4me3 (panel c), H3K4me1 (panel d), H3K27ac (panel e), and H3K27me3 (panel f) were analyzed regarding their association with CpG island (CGI) promoters vs. non-CGI promoters (CGIs were determined in the ±1 kb window of TSSs). The signals relative to the TSS are summarized and the time points compared as indicated. (g) ChIP-qPCR experiments of H3K4me3 measured upstream (5’) or downstream (3’) of the TSS of *Rab8a*. Mean values ±SD of 5-8 experiments are shown.

Our previous studies indicated that CGIs are particularly sensitive to altered KMT2 activities^39^, consistent with previous suggestions^11,16,49^. However, it was unclear whether this was a secondary effect of the slow loss of Ash2l upon recombination. It has been suggested that roughly 70% of vertebrate promoters are characterized by a CGI^50,51^. Of the 25431 H3K4me3 peaks, 18791 are promoter-associated. Of these 79% are characterized by a CGI. To assess whether changes in histone marks are preferentially at CGI promoters upon short-term regulation of FKBP-HA2-ASH2L, we compared the changes in the four histone marks H3K4me3, H3K4me1, H3K27ac and H3K27me3 regarding their association with CGI positive and negative promoters. As soon as significant changes in signal intensities were detected, the majority of these changes were associated with CGI promoters (Fig. 5c-f). Of the promoters with a decrease of H3K4me3 signal in the ±3 kb window of TSSs, 79% were categorized as CGI promoters both after 8 and 16 hours (Fig. 5c). For those that gained H3K4me1 signals, the proportion of CGI promoters was slightly higher, 84% and 82% after 8 and 16 hours, respectively (Fig. 5d). Similar numbers were obtained for promoters that lost H3K27ac signals, 85% after 8 and 16 hours (Fig. 5e). The preference for CGI promoters was higher when H3K27me3 gains were analyzed with 90% and 93% after 8 and 16 hours, respectively, although the numbers are small and therefore the conclusion may not be robust (Fig 5f). Thus, short-term regulation of FKBP-HA2-ASH2L did not reveal a preference for CGI promoters regarding H3K4me3. Nevertheless, the consequences, i.e. the measured increase in H3K4me1, decrease in H3K27ac, and increase in H3K27me3, were slightly higher for CGI promoters than predicted. In other words, promoters that do not possess a CGI seem slightly less responsive to the downstream consequences of FKBP-HA2-ASH2L and H3K4me3 loss, at least at these early time points. Also, similar to the findings upon *Ash2l* KO^39^, the fold decrease was higher upstream of the TSS as exemplified for the *Rab8a* promoter (Fig. 5b and g). Together, this suggests that CGI promoters are only slightly more sensitive to KMT2 complexes upon short-term FKBP-HA2-ASH2L loss, unlike what we observed previously^39^.

### Loss of ASH2L alters chromatin accessibility

The loss of H3K4me3 at promoters and the decrease in gene expression of many genes suggested that promoters may become less accessible. ATAC-seq experiments revealed changes in accessibility that reached 13% of the total accessible sites (153924) across all samples (Fig. 6a, Supplementary Table S8a and b). The number of sites with increased and decreased accessibility was roughly equal. Of these, a proportion was associated with the ±1 kb window surrounding TSSs, but most of the changes were observed in regions that were not associated with promoters, both for regions that gained and lost signals (Fig. 6b). For promoter-associated sites, a slight preference for CGI promoters was observed comprising roughly 82% after 24 hours (Fig. 6c), and thus being comparable to the observations made for H3K4me3, H3K4me1 and H3K27ac (Fig. 5). Although the numbers were small, CGI promoters with gained ATAC signals were underrepresented with 25% (Fig. 6c). Overall, the accessibility at promoters decreased over time (Fig. 6b-d and Supplementary Fig. S4a-c), exemplified at the *Egr1* locus (Supplementary Fig. S2e). The nucleosome-depleted regions, ATAC fragments that are smaller than 120 bp, and well-positioned nucleosomes, ATAC fragments between 130 and 200 bp, at promoters decreased (Fig. 6d and Supplementary Fig. S4b-c). This is consistent with an increase in histone H3 binding at selected promoters (Supplementary Fig. S4d and e), similar to our previous observations in the *Ash2l* KO system^39^. Together, these findings support the notion that chromatin compaction increased. One expectation was that the precise positioning of a nucleosome upstream of the TSS would increase. However, both the nucleosome-free and mono-nucleosome signals decreased, suggesting that the increased protection is not due to precise positioning of nucleosomes. Rather, chromatin seems to appear less ordered. These changes were slow with little difference when the promoters of the fast and slow classes were compared (Fig. 6e) or the groups with high, medium and low H3K4me3 (Supplementary Fig. S4f). This suggests that they are not a direct consequence of ASH2L loss but that they occur downstream of altered histone modifications, including loss of H3K4me3, gain of H3K4me1 and loss of H3K27ac.

**Figure 6.**
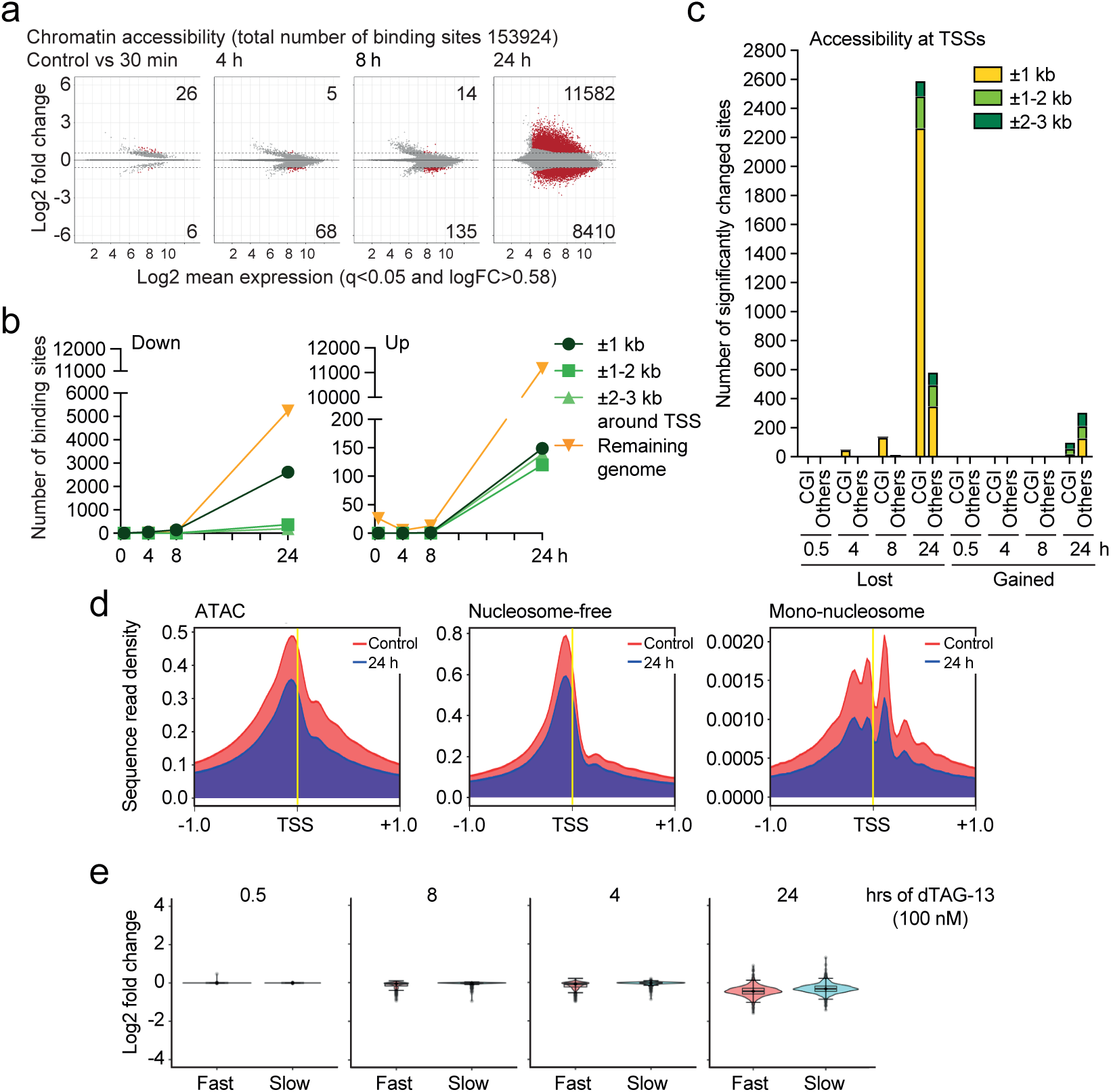
Loss of ASH2L reduces accessibility and decreases regularity at promoters. (a) ATAC-seq analyses were performed on NG3 cells treated with or without dTAG-13 (100 nM) for the indicated times. Displayed are MA plots. Given are the total number of sites identified across all samples and the sites that were significantly more or less accessible (red dots: q<0.05 and logFC>0.58) upon loss of FKBP-HA2-ASH2L. (b) Summary graphs that detail the annotation of significantly changed sites (q<0.05 and logFC>0.58) associated with transcriptional start sites (TSSs) compared to the remaining genome. (c) Promoter-associated ATAC-seq sites with significantly increased or decreased signal intensity were assigned to promoters with or without CpG islands (CGIs were determined in the ±1 kb window of TSSs). (d) Plot profiles generated using DeepTools showing the average normalized signals of the ATAC-seq data are plotted ±1 kb of TSSs of all annotated transcripts in mm10 (normalized using counts per Million). Nucleosome-free and mono-nucleosomes refer to ATAC-seq fragments that are smaller than 120 bp and between 130 and 200 bp, respectively. (e) Promoters with significant decrease in H3K4me3 signals within 2 hours (499, fast) or after 16 hours (788, slow, excluding promoters that lost H3K4me3 signals after 30 min, 2, 4 or 8 hours) (q<0.05; logFC>0.58). A window of ±1 kb around TSSs was considered. The log2 fold changes of the ATAC-seq signals at the promoters of these two classes are displayed.

### Reduced chromatin accessibility at enhancers

To define enhancers, we combined the ATAC, H3K4me1 and H3K27ac signals that were not associated with promoters, i.e., excluding signals associated with a window of ±1 kb of TSSs. This analysis resulted in 32963 putative enhancers regions (Fig. 7a-c, Supplementary Table S9). These regions were also positive for FKBP-HA2-ASH2L, which was broadly reduced upon dTAG-13 treatment (Fig. 7a). Moreover, a substantial fraction of the lost H3K4me1 and H3K27ac signals were associated with enhancers at the 16 hour time point, while only few signal gains were measured (Fig. 7b). The majority of altered ATAC-seq signals (24 h time point) associated with the putative enhancer fraction were reduced, unlike the sites in the remaining genome that increased (Fig. 7b and c). It is important to note that the majority of the enhancers defined by ATAC, H3K4me1 and H3K27ac signals were not affected upon loss of FKBP-HA2-ASH2L when ATAC, H3K4me1 and H3K27ac signals were analyzed. The relatively little changes in H3K4me1 suggests that demethylation at enhancers is slow, similar to the observation at promoters. The H3K4me3 signals were low at enhancers and further decreased over time, while the H3K27me3 signals were very low (Fig. 7a). These findings suggest that the two characteristic histone marks at enhancers, H3K4me1 and H3K27ac, were predominantly downregulated upon loss of FKBP-HA2-ASH2L and that accessibility decreased.

**Figure 7.**
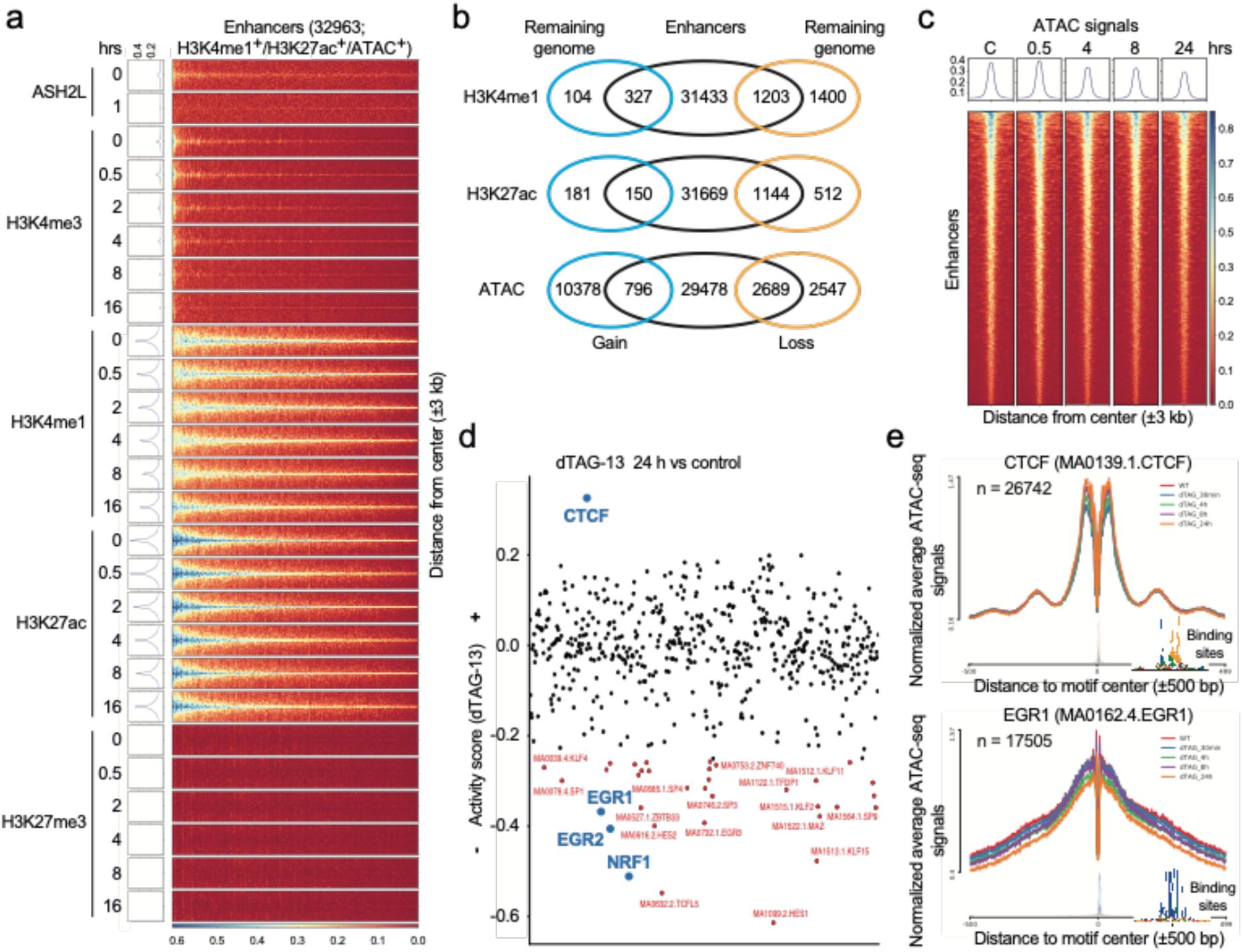
Minor alterations at enhancer upon loss of ASH2L. (a) Enhancers were defined by positivity for H3K4me1, H3K27ac and ATAC signals excluding signals that overlap with ±1 kb of transcriptional start sites (TSSs) of the known mm10 promoters. Heatmaps and plot profiles generated using DeepTools show the indicated ChIP-seq signals (normalized using counts per Million) of Ash2l and the indicated histone marks in response to dTAG-13 treatment. The signals at ±3 kb of the center of the 32963 identified enhancers are displayed. (b) Summarized are the overlaps between significantly changed H3K4me1, H3K27ac, and ATAC sites (q<0.05 and logFC>0.58; the 16 and 24 hour time points for H3K4me1 and H3K27ac and for ATAC-seq, respectively) that are intergenic (more than 3 kb from TSSs of all transcripts in mm10) and the list of enhancers identified as in panel a. Remaining genome refers to the intergenic sites that are not showing overlap with the enhancers as defined here. (c) Heatmaps and plot profiles as defined in panel (a) for ATAC-seq signals centered at enhancers (±3 kb). (d) Analysis of transcription factor (TF) footprinting and their differential activity within the ATAC-seq dataset were conducted using the RGT-HINT tool. TFs highlighted in red are those that have over 1000 identified binding sites and showed significant change (q < 0.05) upon addition of dTAG-13 at 24 hours. (e) Examples of line plots for CTCF (enhanced) and EGR1 (reduced) upon loss of FKBP-HA2-ASH2L. The inserts show the consensus sequence. The indicated numbers (n) are the identified binding sites for the respective TF.

The altered accessibility of chromatin, both at promoters and enhancers, suggested that binding sites of transcription factors might be affected. Therefore, the ATAC fragments were screened for transcription factor (TF) binding sites. Because little effects were observed at early time points (Fig. 6a), we focused on the treatment with dTAG-13 for 24 h. We considered sites that significantly changed upon loss of FKBP-HA2-ASH2L (p < 0.05) and that were represented by at least 1000 binding sites (Supplementary Table S10). With these criteria, more than 30 motifs showed reduced accessibility, including those for EGR1, EGR2 and NRF1 (Fig. 7d). In contrast, only CTCF consensus binding sites showed enhanced binding activity, as previously observed in the KO system^39^. The alterations were time-dependent, exemplified for CTCF and Egr1, but rather small at early time points (Fig. 7e). Thus, the changes in histone marks and chromatin accessibility reduced occupancy of many transcription factor binding sites, compatible with overall decreased accessibility and altered gene transcription.

### Transcriptional rate upon loss of ASH2L

The loss of positive histone marks as well as reduced accessibility at both promoters and enhancers suggested that this affects gene transcription. Indeed, many genes were deregulated in response to FKBP-HA2-ASH2L loss after 24 hours (Fig. 2). To address whether these observations resulted in altered transcription at early time points, we measured newly synthesized RNA by incubating the cells with 5-ethynyl uridine from 3-5 and 7-9 hrs after addition of dTAG-13. The modified RNAs were biotinylated (Click-It), poly-A selected, and sequenced. For control, the expression of FKBP-HA2-ASH2L was visualized in parallel samples (Fig. 8a insert). The analysis revealed that few genes were altered (Fig. 8a, Supplementary Table S11). Thus, at these early time points the loss of FKBP-HA2-ASH2L and the decrease in H3K4me3 at promoters did not have an immediate consequence on gene transcription. We conclude that the loss of FKBP-HA2-ASH2L per se was not sufficient to alter gene expression. It is likely that at later time points effects will become apparent, with the drawback that secondary effects may influence the outcome. To address whether ASH2L can activate gene transcription, we employed a dCAS9 system with sgRNAs containing MS2 binding sites targeting the *HBG1* gene locus^52,53^. In HEK293 cells MS2-ASH2L was sufficient to activate the expression of the chromatin-embedded *HBG1* locus (Fig. 8b). ASH2L interacts with the KMT2 complex core components RBBP5 and DPY30 through the SPRY and the SDI domains, respectively^27,54^. These interactions are necessary for efficient methyltransferase activity and for positioning the complexes to specific sites in chromatin. Mutants in ASH2L that do not interact with RBBP5 and DPY30, MS2-ASH2LΔSPRY and MS2-ASH2LΔSDI, respectively, were unable to induce gene expression (Fig. 8b). Thus, positioning ASH2L, dependent on its ability to interact with its direct partners, is sufficient to induce gene expression, supporting the conclusion that H3K4me3 contributes to transcription^55^.

**Figure 8.**
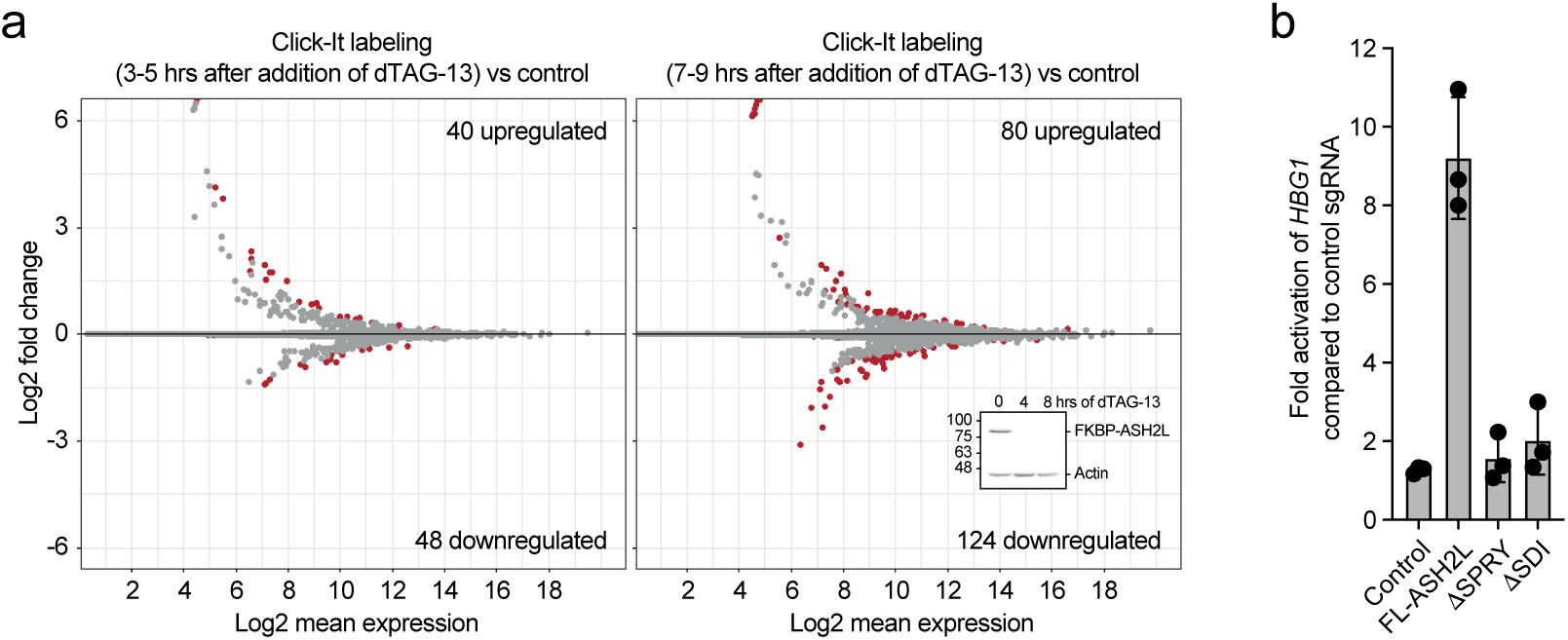
Promoter recruitment of ASH2L stimulates gene expression. (a) Cells were labeled with EdU after the addition of dTAG-13 as indicated. The modified RNA was isolated and analyzed by 3’ end sequencing. MA plots are displayed. Given are the total number of significantly upregulated and downregulated transcripts (red dots; q<0.05). The data summarize two replicates normalized using ERCC spike-in RNA. The insert shows FKBP-HA2-ASH2L protein expression in control cells and cells treated with dTAG-13 for the indicated times. (b) Plasmids expressing dCAS9 and 3 different sgRNAs containing MS2 binding loops were transfected into HEK293 cells. The sgRNAs target the *HBG1* locus. For control, sgRNAs with no specificity for *HBG1* were used. RT-qPCR analysis of *HBG1* RNA was measured, compared to *GUS* RNA and ratios between the measurements of specific sgRNAs vs control sgRNAs are displayed. Mean values ± SD of three experiments.

## Discussion

PROTAC systems have been established for all four WRAD core complex components, including our study^55–57^. This has allowed studying COMPASS/KMT2 complexes and their activities in unprecedented detail. This is important because of the central and broad activity of these complexes that modify both promoters and enhancers with H3K4me3 and H3K4me1, respectively, which are closely linked to gene expression. Obviously, loss of KMT2 complexes and as a consequence altered modifications of H3K4 methylation have very broad effects on gene transcription and RNA expression. Some of these effects are direct but others, probably the majority, are indirect and thus the consequences on biological processes are extensive. The rapid degradation of WDR5, RBBP5, ASH2L and DPY30, in all cases the proteins were lost very rapidly (Figs. 1 and S1; (Refs^55–57^)), allows now to address the direct consequences more specifically. In all cases, this resulted in a decrease of H3K4me3 and less prominent H3K4me1, and broadly altered gene transcription. Because the stability of the H3K4 methylation is different when trimethylation is compared to mono-methylation, the demethylases for these modifications appear to have distinct activities and possibly also distinct mechanisms how they are targeted to chromatin sites. Both knockout and inhibitor studies support an important role for KDM5 family members^55,57^, which are thought to be major H3K4me3 and me2 demethylases^8^. Thus, the dynamics of H3K4 methylation/demethylation is complex and most likely promoter specific.

The dynamics upon loss of WRAD components seem to be different between experimental systems. For example, the effects on H3K4 methylation were much faster in one system of mESCs^55^, but considerably slower in another^57^, the latter being comparable to our iMEF cells. A major difference between mESCs employed in Wang et al. (Ref^55^) and our iMEF cells is that the former grow considerably faster, roughly 12 vs 24 hrs doubling time. This rapid cell cycle progression may require that adjustments to H3K4me3 levels at promoters are much faster with more rapid turnover rates. Indeed, both global as well as promoter-associated H3K4me3 is more stable in our cells (Figs. 1 and 2) than in the mESCs^55^. This is also consistent with the changes in gene transcription that are much faster in mESCs compared to the iMEF cells, despite the three WRAD components ASH2L, DPY30, and RBBP5 being equally fast degraded upon addition of a PROTAC. In the slower mESC system^57^, the effects have a comparable dynamic as observed in our iMEF cells. Together, these findings provide an explanation as to why the timing of downstream effects may not be easily comparable (see also below).

ASH2L is necessary for proliferation, a conclusion that is supported by the findings in the PROTAC system shown here (Fig. 1), and by our previous findings upon knockout of *Ash2l* in MEF cells^40^. Moreover, the knockout of *Ash2l* both in the liver and in the hematopoietic system inhibits proliferation^33,58^. Similarly, the knockout of *Dpy30* prevents proliferation of hematopoietic cells^34^. The knockout of *Wdr5* in mouse embryonic stem cells (mESCs) blocks the ability of these cells to form colonies^59^. Knockdown studies in tumor cells revealed that the different WRAD subunits are important for proliferation to various degrees^10^. In mESCs, which proliferate very fast, the efficient knockdown using PROTAC systems for Rbbp5 and Dpy30 increased duplication time from about 12 to 24 hrs^55^. Thus, both the Rbbp5 and the Dpy30 knockdown cells still proliferate exponentially at least until day 6. Interestingly, the colony formation assays were more decisive, possibly because this is more stressful for cells^55^. Thus, all WRAD subunits are necessary for proliferation to various degrees in the different cell types that have been analyzed. Moreover, the heterogenous effects observed at enhancer and promoters and when RNA is analyzed support the observation that upon loss of ASH2L cells arrest throughout the cell cycle. We interpret this observation to the effect that many different key factors responsible for processes such as cell cycle progression and replication may become limiting in individual cells. As a result, cells arrest in different phases of the cell cycle and are unable to continue into the next cell cycle phase in the absence of ASH2L and most likely because of reduced capacity to activate gene transcription. This may prevent the accumulation of cells in one defined cell cycle phase such as G0/G1. Single cell analyses will be required to address this in more detail.

The pattern of H3K4me3 in control cells and of H3K4me1 in cells after loss of FKBP-HA2-ASH2L is similar around TSSs (Figs. 3 and 5). The 5’ signals are typically weaker than the 3’ signals, while the immediate vicinity of the TSS shows little signal. This latter region is particularly accessible in the ATAC experiments and is the region that binds ASH2L (Fig. 5). Thus, it appears that KMT2 complexes interact with the nucleosome-free region, possibly by binding to the RNAPII through WDR82^60–62^, or recruitment of KMT2 complexes by other means, including sequence-specific transcription factors, that connect to the RNAPII complex bound at the TSS^27^. This region loses accessibility upon FKBP-HA2-ASH2L loss and nucleosomes are less well-positioned (Fig. 6 and Supplementary Fig. S4). Thus, loss of accessibility is not correlated with a stabilization of a well-positioned mono-nucleosome upstream of TSSs. In this respect it is of note that Set1A and B are located mainly downstream of the TSS^62^. Of note, the pattern of Set1A and B binding closely reflects the distribution of H3K4me3 at promoters, while the ASH2L signals are not biphasic (Fig. 5). It will be interesting to see whether other KMT2 enzymes are similarly positioned.

The lack of altered H3K27me3 at promoters was somewhat unexpected as H3K4me3 has been suggested to interfere with Polycomb repressive complex 2 (PRC2), the complex that methylates H3K27. This interference can be due to competing interactions of COMPASS with PRC2 with histone tails and/or by direct inhibition of PRC2 by H3K4me3^63–65^. Whether the remaining H3K4me3 and/or H3K27ac marks are sufficient to prevent H3K27me3 remains open. It is possible that relevant thresholds of these modifications have not been reached for efficient PRC2 recruitment. Alternatively, loss of KMT2 complexes and the decrease of H3K4me3 is not sufficient to promote PRC2 recruitment and activity, for example, because positive signals are lacking. PRC2 recruitment is complex, a major determinant being a high CpG content as in CGIs, but other mechanisms have also been discussed^66,67^. Further studies need to address more directly whether loss of ASH2L alters the binding of PRC2 complexes.

The recent findings suggest that RNAPII is affected by H3K4 methylation and KMT2 complexes. An important early suggestion was that H3K4me3 helps recruit RNAPII complexes as the TFIID subcomplex can directly read this histone mark^68,69^. However, in the PROTAC-mediated loss of RBBP5 and DPY30, no effect on TFIID recruitment was observed^57^. Additional mechanisms have been described, including binding by sequence-specific transcription factors and binding to chromatin^27^. Moreover, interactions with CGI promoters have been documented^11^, which is particularly relevant for SET1A and B^62^. Together, these findings provide further evidence of the complexity of the system, with multiple mechanisms being relevant for the interplay with RNAPII complexes. The recent findings suggest that direct recruitment of RNAPII complexes seems independent of the presence of COMPASS/KMT2 complexes^55,57,62^, consistent with the observation that transcription rate is not affected early after loss of ASH2L (Fig. 8). This is also in agreement with RNAPII loading being unaffected upon loss of RBBP5 early but not late after activation of protein degradation^57^. Nevertheless, the role of H3K4me3 is still not fully understood as even small amounts of this histone mark may be sufficient to control RNAPII loading. Unlike loading, COMPASS complexes regulate pausing of RNAPII and premature termination, although the findings are not fully consistent at present^55,57,62.^

The recent findings discussed above strengthen the correlation between recruitment of KMT2 complexes to promoters, the subsequent H3K4me3, and as a consequence effects on gene transcription. This is further supported by positioning the ASH2L protein through dCAS9 to a specific, endogenous promoter. In this setting, ASH2L is sufficient to activate expression of *HBG1* (Fig. 8). This requires the SPRY domain, which interacts with RBBP5, and the SDI domain, which binds DPY30. Thus, it appears that at least the other WRAD complex subunits, but most likely also one of the KMT2 catalytic subunits, need to be present for activating gene expression.

In summary, our study on the PROTAC-dependent, rapid degradation of ASH2L, provides evidence for a hierarchical alteration of histone marks. While reduction of H3K4me3 occurs first, gain and loss of H3K4me1 at promoters and enhancers, respectively, follows, succeeded by a decrease of H3K27ac at both promoters and enhancers. Similarly, a reduction in chromatin accessibility is occurring late, resulting in less structured nucleosomal organization at promoters, while H3K27me3 is only minimally affected. Thus, also the initial event, the degradation of ASH2L is fast, subsequent chromatin-associated steps as well as regulation of gene expression are rather slow, which argues for a strong buffering effect built into the system. This might be due to additional histone marks, which are manifold and most of them only poorly understood mechanistically^5,70,71^. Also, thresholds of distinct histone modifications might influence RNAPII loading and activity. Thus, PROTOAC systems as used here and published by others while our work was in progress^55,57,62^, support a determining role of H3K4 methylation in controlling gene transcription and will allow further detailed evaluation to define functions of COMPASS/KMT2 complexes.

## Material and Methods

### Protein analyses

SDS-PAGE and Western blotting have been performed identically to the procedure described previously and the antibodies used have been listed^40^.

### RNA isolation and cDNA synthesis

RNA purification from cells was performed using the HighPure RNA Isolation Kit (Roche, 11826665001). The RNA was transcribed into cDNA by using the QuantiTect Reverse Transcription Kit (Qiagen, 205314). For *Il7* and *Egr1* QuantiTect primers (QT00101318 and QT00265846, respectively, Qiagen) were used for quantiative PCR. Primers for *Nrf1* were: forward, GAGAATGTGGTGCGAAAGT; reverse, GCTCTGAATTAACCTCCTGTG; for *Ppbp*: forward, ACCATCTCTGGAATCCCATTCA; reverse, GTCCATTCTTCAGTGTGGCTATC.

### Cloning and Vector Design

The human ASH2L sequence was introduced to the lentiviral vector pLEX305-NdTAG (Addgene#91797) using a Gateway LR reaction (Invitrogen). The LR reactions were performed overnight at 25°C in a final volume of 10 μl according to the manufacturer’s instruction and used to transform competent Stbl3 bacteria. After plasmid preparation the integrity of the Gateway expression constructs was controlled by restriction digest.

### Cell culture and lentiviral transduction

HEK293T and immortalized mouse embryonic fibroblast (iMEF^40^) cells were cultured in DMEM supplemented with 10% (v/v) FBS and 1% P/S (v/v) at 37 °C in a humidified incubator at 5% CO2.

For lentiviral vector production, a third-generation packaging system was transfected into HEK293T cells, including three helper plasmids (pMDLg/pRRE, pRSV-Rev, pCMV-VSV-G), pLEX-305-N-dTAG-hASH2L expressing FKBP-HA2-ASH2L, and pH2B-YFP to estimate transfection efficiency. After 48 hours, the cell culture supernatant was collected and passed through a 0.45 µm PVDF filter. iMEF cells were seeded at a confluency of ∼40% and incubated with lentivirus-containing supernatant, diluted 1:2 with fresh medium, on the next day. Polybrene (8 µg/mL) was included in the infection protocol. The medium was changed after 6-8 hrs and puromycin (8 µg/mL) was added the following day. In order to eliminate the endogenous *Ash2l*, transduced cells were treated with 5 nM of 4-hydroxytamoxifen for 3 days to excise exon 4 of the endogenous *Ash2l* loci. Individual clones were then established using limiting dilution and expanded.

### Cell Proliferation Assay

For cell proliferation assays, three biological replicates, each in triplicates, were performed. Five x 10^4^ cells were seeded in a 6-well plate and dTAG-13 treatment (100 nM) immediately. Cells were counted at days as indicated in the figure. Before cells reached confluency, they were split at a ratio of 1:2–1:3.

### Cell cycle analysis

Cells were harvested by trypsinization, collected by centrifugation and washed in ice-cold phosphate-buffered saline (PBS). The cells were then fixed using 4% paraformaldehyde in PBS, and stained with Vybrant™ DyeCycle™ Violet Stain (Invitrogen™, #V35003) at the final concentration of 5 µM at 37 °C for 30 min. Violet signal was acquired in the Pacific Blue channel in a linear mode with low speed. The percentage of the cell population in distinct phases of the cell cycle was determined using a manual gating method. G1 gate width was considered the same as G2 gate width (unconstrained), and the area under the curve was calculated as cell percentage.

### EdU Incorporation Assay

Click-iT™ EdU Alexa Fluor™ 488 (AF488) Flow Cytometry Assay kit (Invitrogen) was used to analyze DNA synthesis essentially according to the manufacturer’s instruction. The labeling with AF488-Conjugated EdU (5-ethynyl-2’-deoxyuridine) was done at a final concentration of 10 µM for 3 hours prior to harvesting. The DNA was stained with Vybrant DyeCycle Violet. Violet and AF488 signals were acquired in the Pacific Blue (Lin), and FITC (Log) channels, respectively. The percentage of FITC-positive cells was determined using manual gating with a threshold set above the FITC signal from non-EdU treated samples.

### RNA-Seq

RNA was isolated as described above. For quality control, samples were analyzed using an RNA ScreenTape (Agilent, 5067-5576) with a TapeStation-device (Agilent). For internal control, ERCC-RNA-spike-in (ThermoFisher, 4456740) was added to every sample. For library generation the Collibri 3’-mRNA Prep Kit (ThermoFisher, A38110024) was used. Sequencing was performed on a NextSeq500/550 platform (Illumina) based on a Mid Output Kit 2.5 (Illumina, 20224904) using 75 cycles and single-end reads. Quality control, library preparation and sequencing were executed by the Genomics Core Facility of the Interdisciplinary Center for Clinical Research (IZKF) of the Medicine Faculty at RWTH Aachen University.

### Chromatin-Immunoprecipitation (ChIP)

ChIP experiments were carried out using the ChIP-IT High Sensitivity® kit (ActiveMotif) essentially according to the manufacturer’s instruction. Modifications were as follows. For nuclei isolation, 60-70 strokes were applied using a 5 mL glass dounce homogenizer with a tight (B) pestle. The amount of chromatin used per IP was 30 µg and 100 µg for histone marks and ASH2L, respectively. Chromatin shearing was conducted using the Bioruptor® Pico sonication device. Sonication was performed in 300 µL aliquots in 1.5 mL Bioruptor® Pico microtube for 4-5 rounds of 10 cycles (each cycle was 30s sonication/30s pause) until the majority of chromatin fragments were sheared down to ∼200 and to ∼500 bp (for ChIP-seq and ChIP-RT-qPCR, respectively). Input DNA was precipitated by adding 2 µL of carrier (provided by the kit) and 2 µL glycogen (20 mg/mL). For sequencing, the concentration of samples was measured using the Quantus™ Fluorometer. Sample quality control/fragment size distribution was assessed using the Bioanalyzer system (Agilent). Samples were then indexed and adaptor-ligated using NEBNEXT Ultra II DNA Library Preparation Kit (NEB) according to the manufacturer’s instructions. For all histone marks (except H3K27me3, known as broad histone mark) a ∼350 bp (200 bp chromatin fragment size + 120 bp two adaptors size) size selection step was conducted prior to PCR library amplification. Sequencing was performed on a NextSeq 550 (Illumina) system using a NextSeq 500/550 High Output v2.5 (75 Cycles) cartridge (Single-End). The number of samples per cartridge was arranged in a way to provide a minimum of 40 – 50 M raw reads per ChIP sample.

### ATAC seq

Chromatin accessibility was performed as described^39,72–74^. In brief, cells were treated with dTAG-13 or EtOH for control, collected and washed in PBS. Subsequently, ATAC lysis buffer (10 mM Tris-HCl, pH 7.4, 10 mM NaCl, 3 mM MgCl2, 0.1% NP-40, 0.1% Tween-20, 0.01% Digitonin) was added. Nuclei were pelleted and incubated in transposition mix (1x TD (Illumina, 20034210), 7.5 μl per sample TDE1 (Illumina, 20034210), 0.1% Tween-20, 0.01% Digitonin, in PBS) in a final volume of 50 μl per sample at 37°C for 1 h. Tagmented fragments were purified using the MinElute PCR purification kit (Qiagen, 28004). The transposed DNA fragments were amplified using 25 µM Nextera i5 and i7 barcoded primers in NEB Next Ultra II Q5 Master Mix (New England Labs, M0544). DNA purification was performed using AMPureXP magnetic beads (Beckman Coulter, A63880) in 96-well plates. The beads were washed with 200 µl of 85% ethanol. The DNA was eluted in water and stored in DNA low-binding tubes.

### Click-It-seq

Newly synthesized transcripts were analyzed using the Click-iT™ Nascent RNA Capture Kit (Invitrogen). The cells were incubated with 0.2 mM EdU for 2 hrs. The sample processing was conducted according to the manufacturer’s instructions. After the final RNA pull-down, biotinylated RNAs were separated from the beads in TRIzol^TM^ reagent. Sample quality control/fragment size distribution was assessed using the Bioanalyzer system (Agilent). The concentration of samples was measured using the Quantus™ Fluorometer. Isolated RNA samples with an RNA Integrity Number (RIN) higher than 9 were validated for further sequencing analysis. Construction of cDNA libraries from RNA samples was done using the Collibri™ 3′ mRNA Library Prep Kit (Invitrogen). Samples were sequenced on a NextSeq 550 (Illumina) system using a NextSeq 500/550 Mid Output v2.5 (75 Cycles) cartridge (Single-End). The number of samples per cartridge was arranged in a way to provide a minimum of 20 M raw reads per 3’mRNAseq sample.

### dCAS9-mediated gene expression

Vectors for dCAS9 and sgRNAs containing MS2 binding sites are based on previous reports^53,75^. Sequences encoding MS2 and 3 Flag-tags were obtained as gene blocks (Integrated DNA Technologies (IDT)) and cloned into a CMV-driven vector containing a Gateway cloning cassette. Sequences for ASH2L and mutants were recombined into this vector. HEK293 cells were co-transfected with plasmids expressing dCAS9, sgRNAs targeting the *HBG1* promoter^53^, and MS2 and MS2-ASH2L fusion proteins expressing vectors. RNA was harvested and reverse transcribed as described above. *HBG1* expression was analyzed using RT-qPCR assays with QuantiTect primers. For control, *β-glucuronidase* was measured (forward: CTCATTTGGAATTTTGCCGATT; reverse: CCGAGTGAAGATCCCCTTTTTA).

### Bioinformatics

The raw reads of all sequencing experiments (ChIP-seq, ATAC-seq, RNA-seq and Click-it) were trimmed using Trim_Galore (https://www.bioinformatics.babraham.ac.uk/projects/trim_galore/). The reference genome used here was the mouse reference genome mm10 (GRCm38.p6; http://ftp.ebi.ac.uk/pub/databases/gencode/Gencode_mouse/release_M25/). BWA was used to align the trimmed reads from ChIP-seq and ATAC-seq to mm10 (Ref^76^). The Trimmed reads from RNA-seq and Click-it were aligned to mm10 using STAR^77^. Samtools were used to generate the sorted Bam files after filtering the unusable reads as recommended by Encode^76^. Quality controls for both ChIP-seq and ATAC-seq were done as recommended by Encode. Narrow peaks (ChIP-seq (H3K4me3, H3K27ac, ASH2L) and ATAC-seq) and broad peaks (ChIP-seq (H3K4me1, H3K27me3)) were called using Macs^78^. In RNA-seq and Click-it, the reads aligned to the mm10 genome were assigned to the known annotated mm10 genes using featureCounts^79^. The counts obtained from featureCounts were normalized using the External RNA Control Consortium (ERCC, Invitrogen) spike-in. DeepTools were used to generate both heatmaps and plotprofiles showing the normalized signal (normalized using CPM (count per Million)) for the ChIP-seq or ATAC-seq experiments^80^. This was centered either at TSS of all annotated transcripts in mm10 or at enhancers we identified during this work. Enhancers were defined by positivity for H3K4me1, H3K27ac and ATAC signal in the control samples excluding the overlap with -+1kb of TSS of the known mm10 promoters. MA plots were generated using the ggmaplot function of ggplot2 package in Rstudio^81^. The apeglm method for log2 fold change shrinkage was applied for the MA plots^82^. The sampleDistMatrix function of pheatmap package in Rstudio was used to generate the heatmaps (Raivo Kolde (2019). pheatmap: Pretty Heatmaps. R package version 1.0.12. http://CRAN.R-project.org/package=pheatmap). The promoters significantly lost H3K4me3 (q<0,05; logFC>0.58) were categorized as fast and slow as follows: fast (499 promoters) are all promoters that showed a significant loss of H3K4me3 signals after 2 h, slow (788 promoters) are all promoters that showed a significant loss of H3K4me3 signals only after 16 hours (excluding promoters significantly losing H3K4me3 signal after 30 min, 2 h, 4 h or 8 h). In addition, all identified H3K4me3 across all samples (25431 sites) were divided into three equal categories according to the signal intensity: high, medium and low (8477 each). The overlap between different comparisons in the same sequencing experiment and between different experiments were done using BEDTools^83^. ChIP-seq samples were normalized to the lowest coverage in each experiment. In ATAC-seq, the data was normalized using Deseq2^84^. Differential analysis for each ChIP-seq and ATAC-seq, RNA-seq and Click-it experiments was carried out using Deseq2 to identify the significantly changed sites. These sites were then annotated using Homer (http://homer.ucsd.edu/homer/ngs/annotation.html). The information after annotation (distance to the nearest promoter provided by Homer) was used to identify the distance to TSS. Coordinates of CGI in mm10 were obtained from UCSC (https://hgdownload.soe.ucsc.edu/goldenPath/mm10/database/cpgIslandExt.txt.gz). The replicates were merged using Picard MergeSamFiles (https://github.com/broadinstitute/picard/blob/master/src/main/java/picard/sam/MergeSamFiles.java). In ATAC-seq, transcription factor (TF)-footprinting analysis was performed using RGT-HINT^73^. Sections of the codes from nf-core were modified and used for ChIP-seq (https://nf-co.re/chipseq/) and ATAC-seq (https://nf-co.re/atacseq) analyses. IGV (Integrative Genomics Viewer, https://software.broadinstitute.org/software/igv/) was used to visualize the normalized BigWig files and to evaluate the results.

All sequencing data are available in NCBI’s Gene Expression Omnibus (GEO; Ref^85^) as SuperSeries under accession number GSE241174. This SuperSeries is composed of the following sub-series: 1. Accession number GSE240987 for RNA-seq. 2. Accession number GSE241001 for ASH2L ChIP-seq. 3.Accession number GSE240994 for H3K4me3 ChIP-seq. 4. Accession number GSE240992 for H3K4me1 ChIP-seq. 5. Accession number GSE240990 for H3K27ac ChIP-seq. 6. Accession numbers GSE240999 and GSE241000 for H3K27me3 ChIP-seq. 7. Accession number GSE241169 for ATAC-seq. 8. Accession number GSE239789 for Click-it-seq.

## Acknowledgments

We thank A. Golzmann, S. Krieg, A. Redecker, and R. Zaja for providing vectors for cloning; J. Franzen, L. Gan, J. Hübner and J. Kuo of the genomics core facility of the Interdisciplinary Center for Clinical Research (IZKF) Aachen, Faculty of Medicine, RWTH Aachen University for sample preparation and sequencing. Simulations were performed with computing resources granted by RWTH Aachen University under project rwth0751 (to M.B.). The work was funded by grants from the German Research Foundation DFG (LU466/17-2 to B.L.) and the START program of the Faculty of Medicine, RWTH Aachen University (129/22) (to M.B.).

## Author contributions

R.S.B., A.T.S., and P.B. performed the wet experiments; M.B. carried out bioinformatic analyses; J.L.F and B.L. developed the project; M.B., R.S.B., A.T.S., J.L.F., and B.L. analyzed data; B.L. wrote the initial version of the paper; all authors read and approved the final manuscript.

## Supplementary information

Supplementary Figures S1-4

### Supplementary Tables

All sequencing data are available in NCBI’s Gene Expression Omnibus as SuperSeries under accession number GSE241174.

**Table S1** List of significantly changed genes after 24h compared to the WT (RNA-seq)

**Table S2a** All identified Ash2l peaks across all the samples

**Table S2b** List of significantly changed Ash2l binding sites after 1h compared to the WT (ChIP-seq Ash2l)

**Table S3a** All identified H3K4me3 peaks across all the samples

**Table S3b** List of significantly changed H3K4me3 binding sites in each timepoint compared to the WT (ChIP-seq H3K4me3)

**Table S4a** All identified H3K4me1 peaks across all the samples

**Table S4b** List of significantly changed H3K4me1 binding sites in each timepoint compared to the WT (ChIP-seq H3K4me1)

**Table S5a** All identified H3K27ac peaks across all the samples

**Table S5b** List of significantly changed H3K27ac binding sites in each timepoint compared to the WT (ChIP-seq H3K27ac)

**Table S6a** All identified H3K27me3 peaks across all the samples (ChIP-seq H3K27me3_First experiment)

**Table S6b** List of significantly changed H3K27me3 binding sites in each timepoint compared to the WT (ChIP-seq H3K27me3_First experiment)

**Table S7a** All identified H3K27me3 peaks across all the samples (ChIP-seq H3K27me3_Second experiment)

**Table S7b** List of significantly changed H3K27me3 binding sites in each timepoint compared to the WT (ChIP-seq H3K27me3_Second experiment)

**Table S8a** All identified peaks across all the samples (ATAC-seq)

**Table S8b** List of sites significantly changed in accessibility in each timepoint compared to the WT (ATAC-seq)

**Table S9** List of all identified putative Enhancers

**Table S10** List of significantly changed TF (ATAC-seq; differential analysis using RGT-Hint)

**Table S11** List of significantly changed genes after 4h or 8h compared to the WT (Nascent RNA; Click-it)

## References

1. Bannister, A. J. & Kouzarides, T. Regulation of chromatin by histone modifications. Cell Research 21, 381–395, doi:10.1038/cr.2011.22 (2011).

2. Hafner, A. & Boettiger, A. The spatial organization of transcriptional control. Nat Rev Genet 24, 53–68, doi:10.1038/s41576-022-00526-0 (2023).

3. da Costa-Nunes, J. A. & Noordermeer, D. TADs: Dynamic structures to create stable regulatory functions. Curr Opin Struct Biol 81, 102622, doi:10.1016/j.sbi.2023.102622 (2023).

4. Zentner, G. E. & Henikoff, S. Regulation of nucleosome dynamics by histone modifications. Nature structural & molecular biology 20, 259–266, doi:10.1038/nsmb.2470 (2013).

5. Morgan, M. A. J. & Shilatifard, A. Reevaluating the roles of histone-modifying enzymes and their associated chromatin modifications in transcriptional regulation. Nat Genet 52, 1271–1281, doi:10.1038/s41588-020-00736-4 (2020).

6. Soto, L. F. et al. Compendium of human transcription factor effector domains. Mol Cell 82, 514–526, doi:10.1016/j.molcel.2021.11.007 (2022).

7. Kim, S. & Wysocka, J. Deciphering the multi-scale, quantitative cis-regulatory code. Mol Cell 83, 373–392, doi:10.1016/j.molcel.2022.12.032 (2023).

8. Pavlenko, E., Ruengeler, T., Engel, P. & Poepsel, S. Functions and Interactions of Mammalian KDM5 Demethylases. Front Genet 13, 906662, doi:10.3389/fgene.2022.906662 (2022).

9. Shilatifard, A. The COMPASS family of histone H3K4 methylases: mechanisms of regulation in development and disease pathogenesis. Annu Rev Biochem 81, 65–95, doi:10.1146/annurev-biochem-051710-134100 (2012).

10. Jiang, H. The complex activities of the SET1/MLL complex core subunits in development and disease. Bba-Gene Regul Mech 1863, doi:10.1016/j.bbagrm.2020.194560 (2020).

11. Hughes, A. L., Kelley, J. R. & Klose, R. J. Understanding the interplay between CpG island-associated gene promoters and H3K4 methylation. Biochim Biophys Acta Gene Regul Mech 1863, 194567, doi:10.1016/j.bbagrm.2020.194567 (2020).

12. Park, S., Kim, G. W., Kwon, S. H. & Lee, J. S. Broad domains of histone H3 lysine 4 trimethylation in transcriptional regulation and disease. FEBS J 287, 2891–2902, doi:10.1111/febs.15219 (2020).

13. Calo, E. & Wysocka, J. Modification of enhancer chromatin: what, how, and why? Mol Cell 49, 825–837, doi:10.1016/j.molcel.2013.01.038 (2013).

14. Catarino, R. R. & Stark, A. Assessing sufficiency and necessity of enhancer activities for gene expression and the mechanisms of transcription activation. Genes Dev 32, 202–223, doi:10.1101/gad.310367.117 (2018).

15. Buecker, C. & Wysocka, J. Enhancers as information integration hubs in development: lessons from genomics. Trends Genet 28, 276–284, doi:10.1016/j.tig.2012.02.008 (2012).

16. Macrae, T. A., Fothergill-Robinson, J. & Ramalho-Santos, M. Regulation, functions and transmission of bivalent chromatin during mammalian development. Nat Rev Mol Cell Biol 24, 6–26, doi:10.1038/s41580-022-00518-2 (2023).

17. Voigt, P., Tee, W. W. & Reinberg, D. A double take on bivalent promoters. Genes Dev 27, 1318–1338, doi:10.1101/gad.219626.113 (2013).

18. Hyun, K., Jeon, J., Park, K. & Kim, J. Writing, erasing and reading histone lysine methylations. Exp Mol Med 49, e324, doi:10.1038/emm.2017.11 (2017).

19. Beacon, T. H. et al. The dynamic broad epigenetic (H3K4me3, H3K27ac) domain as a mark of essential genes. Clin Epigenetics 13, 138, doi:10.1186/s13148-021-01126-1 (2021).

20. Miller, T. et al. COMPASS: a complex of proteins associated with a trithorax-related SET domain protein. Proc Natl Acad Sci U S A 98, 12902–12907 (2001).

21. Couture, J. F. & Skiniotis, G. Assembling a COMPASS. Epigenetics 8, 349–354, doi:10.4161/epi.24177 (2013).

22. Patel, A., Dharmarajan, V., Vought, V. E. & Cosgrove, M. S. On the mechanism of multiple lysine methylation by the human mixed lineage leukemia protein-1 (MLL1) core complex. J Biol Chem 284, 24242–24256, doi:10.1074/jbc.M109.014498 (2009).

23. Southall, S. M., Wong, P. S., Odho, Z., Roe, S. M. & Wilson, J. R. Structural basis for the requirement of additional factors for MLL1 SET domain activity and recognition of epigenetic marks. Mol Cell 33, 181–191, doi:10.1016/j.molcel.2008.12.029 (2009).

24. Avdic, V. et al. Structural and biochemical insights into MLL1 core complex assembly. Structure 19, 101–108, doi:10.1016/j.str.2010.09.022 (2011).

25. Steward, M. M. et al. Molecular regulation of H3K4 trimethylation by ASH2L, a shared subunit of MLL complexes. Nat Struct Mol Biol 13, 852–854, doi:10.1038/nsmb1131 (2006).

26. Dou, Y. et al. Regulation of MLL1 H3K4 methyltransferase activity by its core components. Nat Struct Mol Biol 13, 713–719, doi:10.1038/nsmb1128 (2006).

27. Bochynska, A., Luscher-Firzlaff, J. & Luscher, B. Modes of Interaction of KMT2 Histone H3 Lysine 4 Methyltransferase/COMPASS Complexes with Chromatin. Cells 7, doi:10.3390/cells7030017 (2018).

28. Rao, R. C. & Dou, Y. Hijacked in cancer: the KMT2 (MLL) family of methyltransferases. Nat Rev Cancer 15, 334–346, doi:10.1038/nrc3929 (2015).

29. Poreba, E., Lesniewicz, K. & Durzynska, J. Aberrant Activity of Histone-Lysine N-Methyltransferase 2 (KMT2) Complexes in Oncogenesis. Int J Mol Sci 21, doi:10.3390/ijms21249340 (2020).

30. Deaton, A. M. & Bird, A. CpG islands and the regulation of transcription. Genes Dev 25, 1010–1022, doi:10.1101/gad.2037511 (2011).

31. Stoller, J. Z. et al. Ash2l interacts with Tbx1 and is required during early embryogenesis. Exp Biol Med (Maywood) 235, 569–576, doi:10.1258/ebm.2010.009318 (2010).

32. Adamson, A. L. & Shearn, A. Molecular genetic analysis of Drosophila ash2, a member of the trithorax group required for imaginal disc pattern formation. Genetics 144, 621–633 (1996).

33. Luscher-Firzlaff, J. et al. Hematopoietic stem and progenitor cell proliferation and differentiation requires the trithorax protein Ash2l. Sci Rep 9, 8262, doi:10.1038/s41598-019-44720-3 (2019).

34. Yang, Z. et al. The DPY30 subunit in SET1/MLL complexes regulates the proliferation and differentiation of hematopoietic progenitor cells. Blood 124, 2025–2033, doi:10.1182/blood-2014-01-549220 (2014).

35. Yang, Z., Shah, K., Khodadadi-Jamayran, A. & Jiang, H. Dpy30 is critical for maintaining the identity and function of adult hematopoietic stem cells. J Exp Med 213, 2349–2364, doi:10.1084/jem.20160185 (2016).

36. Luscher-Firzlaff, J. et al. The human trithorax protein hASH2 functions as an oncoprotein. Cancer Res 68, 749–758, doi:10.1158/0008-5472.CAN-07-3158 (2008).

37. Ullius, A. et al. The interaction of MYC with the trithorax protein ASH2L promotes gene transcription by regulating H3K27 modification. Nucleic Acids Res 42, 6901–6920, doi:10.1093/nar/gku312 (2014).

38. Wan, M. et al. The trithorax group protein Ash2l is essential for pluripotency and maintaining open chromatin in embryonic stem cells. J Biol Chem 288, 5039–5048, doi:10.1074/jbc.M112.424515 (2013).

39. Barsoum, M. et al. Loss of the Ash2l subunit of histone H3K4 methyltransferase complexes reduces chromatin accessibility at promoters. Sci Rep 12, 21506, doi:10.1038/s41598-022-25881-0 (2022).

40. Bochynska, A. et al. Induction of senescence upon loss of the Ash2l core subunit of H3K4 methyltransferase complexes. Nucleic Acids Res 50, 7889–7905, doi:10.1093/nar/gkac591 (2022).

41. Nabet, B. et al. The dTAG system for immediate and target-specific protein degradation. Nat Chem Biol 14, 431–441, doi:10.1038/s41589-018-0021-8 (2018).

42. Clackson, T. et al. Redesigning an FKBP-ligand interface to generate chemical dimerizers with novel specificity. Proc Natl Acad Sci U S A 95, 10437–10442, doi:10.1073/pnas.95.18.10437 (1998).

43. Winter, G. E. et al. DRUG DEVELOPMENT. Phthalimide conjugation as a strategy for in vivo target protein degradation. Science 348, 1376–1381, doi:10.1126/science.aab1433 (2015).

44. Martinez-Gamero, C., Malla, S. & Aguilo, F. LSD1: Expanding Functions in Stem Cells and Differentiation. Cells 10, doi:10.3390/cells10113252 (2021).

45. Havis, E. & Duprez, D. EGR1 Transcription Factor is a Multifaceted Regulator of Matrix Production in Tendons and Other Connective Tissues. Int J Mol Sci 21, doi:10.3390/ijms21051664 (2020).

46. Thiel, G. & Cibelli, G. Regulation of life and death by the zinc finger transcription factor Egr-1. J Cell Physiol 193, 287–292, doi:10.1002/jcp.10178 (2002).

47. Bhagwat, A. S. & Vakoc, C. R. A new bump in the epigenetic landscape. Mol Cell 53, 857–858, doi:10.1016/j.molcel.2014.03.001 (2014).

48. Cheng, J. et al. A role for H3K4 monomethylation in gene repression and partitioning of chromatin readers. Mol Cell 53, 979–992, doi:10.1016/j.molcel.2014.02.032 (2014).

49. Piunti, A. & Shilatifard, A. Epigenetic balance of gene expression by Polycomb and COMPASS families. Science 352, aad9780, doi:10.1126/science.aad9780 (2016).

50. Saxonov, S., Berg, P. & Brutlag, D. L. A genome-wide analysis of CpG dinucleotides in the human genome distinguishes two distinct classes of promoters. Proc Natl Acad Sci U S A 103, 1412–1417, doi:10.1073/pnas.0510310103 (2006).

51. Long, H. K. et al. Epigenetic conservation at gene regulatory elements revealed by non-methylated DNA profiling in seven vertebrates. Elife 2, e00348, doi:10.7554/eLife.00348 (2013).

52. Maeder, M. L. et al. CRISPR RNA-guided activation of endogenous human genes. Nat Methods 10, 977–979, doi:10.1038/nmeth.2598 (2013).

53. Konermann, S. et al. Genome-scale transcriptional activation by an engineered CRISPR-Cas9 complex. Nature 517, 583–588, doi:10.1038/nature14136 (2015).

54. Ernst, P. & Vakoc, C. R. WRAD: enabler of the SET1-family of H3K4 methyltransferases. Brief Funct Genomics 11, 217–226, doi:10.1093/bfgp/els017 (2012).

55. Wang, H. et al. H3K4me3 regulates RNA polymerase II promoter-proximal pause-release. Nature 615, 339–348, doi:10.1038/s41586-023-05780-8 (2023).

56. Li, D. et al. Discovery of a dual WDR5 and Ikaros PROTAC degrader as an anti-cancer therapeutic. Oncogene 41, 3328–3340, doi:10.1038/s41388-022-02340-8 (2022).

57. Hu, S. et al. H3K4me2/3 modulate the stability of RNA polymerase II pausing. Cell Res 33, 403–406, doi:10.1038/s41422-023-00794-3 (2023).

58. Liang, W. L. et al. Loss of the epigenetic regulator Ash2l results in desintegration of hepatocytes and liver failure. Int J Clin Exp Patho 9, 5167–5175 (2016).

59. Li, Q. et al. p53 inactivation unmasks histone methylation-independent WDR5 functions that drive self-renewal and differentiation of pluripotent stem cells. Stem Cell Reports 16, 2642–2658, doi:10.1016/j.stemcr.2021.10.002 (2021).

60. Lee, J. H. & Skalnik, D. G. Wdr82 is a C-terminal domain-binding protein that recruits the Setd1A Histone H3-Lys4 methyltransferase complex to transcription start sites of transcribed human genes. Mol Cell Biol 28, 609–618, doi:10.1128/MCB.01356-07 (2008).

61. Ebmeier, C. C. et al. Human TFIIH Kinase CDK7 Regulates Transcription-Associated Chromatin Modifications. Cell Rep 20, 1173–1186, doi:10.1016/j.celrep.2017.07.021 (2017).

62. Hughes, A. L. et al. A CpG island-encoded mechanism protects genes from premature transcription termination. Nat Commun 14, 726, doi:10.1038/s41467-023-36236-2 (2023).

63. Douillet, D. et al. Uncoupling histone H3K4 trimethylation from developmental gene expression via an equilibrium of COMPASS, Polycomb and DNA methylation. Nat Genet 52, 615–625, doi:10.1038/s41588-020-0618-1 (2020).

64. Schmitges, F. W. et al. Histone methylation by PRC2 is inhibited by active chromatin marks. Mol Cell 42, 330–341, doi:10.1016/j.molcel.2011.03.025 (2011).

65. Kasinath, V. et al. JARID2 and AEBP2 regulate PRC2 in the presence of H2AK119ub1 and other histone modifications. Science 371, doi:10.1126/science.abc3393 (2021).

66. Laugesen, A., Hojfeldt, J. W. & Helin, K. Molecular Mechanisms Directing PRC2 Recruitment and H3K27 Methylation. Mol Cell 74, 8–18, doi:10.1016/j.molcel.2019.03.011 (2019).

67. van Kruijsbergen, I., Hontelez, S. & Veenstra, G. J. Recruiting polycomb to chromatin. Int J Biochem Cell Biol 67, 177–187, doi:10.1016/j.biocel.2015.05.006 (2015).

68. Vermeulen, M. et al. Selective Anchoring of TFIID to Nucleosomes by Trimethylation of Histone H3 Lysine 4. Cell 131, 58–69 (2007).

69. Lauberth, S. M. et al. H3K4me3 interactions with TAF3 regulate preinitiation complex assembly and selective gene activation. Cell 152, 1021–1036, doi:10.1016/j.cell.2013.01.052 S0092-8674(13)00144-X [pii] (2013).

70. Huang, H., Sabari, B. R., Garcia, B. A., Allis, C. D. & Zhao, Y. SnapShot: histone modifications. Cell 159, 458–458 e451, doi:10.1016/j.cell.2014.09.037 (2014).

71. Talbert, P. B., Meers, M. P. & Henikoff, S. Old cogs, new tricks: the evolution of gene expression in a chromatin context. Nat Rev Genet 20, 283–297, doi:10.1038/s41576-019-0105-7 (2019).

72. Buenrostro, J. D., Giresi, P. G., Zaba, L. C., Chang, H. Y. & Greenleaf, W. J. Transposition of native chromatin for fast and sensitive epigenomic profiling of open chromatin, DNA-binding proteins and nucleosome position. Nat Methods 10, 1213–1218, doi:10.1038/nmeth.2688 (2013).

73. Li, Z. et al. Identification of transcription factor binding sites using ATAC-seq. Genome Biol 20, 45, doi:10.1186/s13059-019-1642-2 (2019).

74. Brunton, H., Garner, I. M., Bailey, U. M., Upstill-Goddard, R. & Bailey, P. J. Using Chromatin Accessibility to Delineate Therapeutic Subtypes in Pancreatic Cancer Patient-Derived Cell Lines. STAR Protoc 1, 100079, doi:10.1016/j.xpro.2020.100079 (2020).

75. Hilton, I. B. et al. Epigenome editing by a CRISPR-Cas9-based acetyltransferase activates genes from promoters and enhancers. Nat Biotechnol 33, 510–517, doi:10.1038/nbt.3199 (2015).

76. Li, H. et al. The Sequence Alignment/Map format and SAMtools. Bioinformatics 25, 2078–2079, doi:10.1093/bioinformatics/btp352 (2009).

77. Dobin, A. et al. STAR: ultrafast universal RNA-seq aligner. Bioinformatics 29, 15–21, doi:10.1093/bioinformatics/bts635 (2013).

78. Zhang, Y. et al. Model-based analysis of ChIP-Seq (MACS). Genome Biol 9, R137, doi:10.1186/gb-2008-9-9-r137 (2008).

79. Liao, Y., Smyth, G. K. & Shi, W. featureCounts: an efficient general purpose program for assigning sequence reads to genomic features. Bioinformatics 30, 923–930, doi:10.1093/bioinformatics/btt656 (2014).

80. Ramirez, F., Dundar, F., Diehl, S., Gruning, B. A. & Manke, T. deepTools: a flexible platform for exploring deep-sequencing data. Nucleic Acids Res 42, W187–191, doi:10.1093/nar/gku365 (2014).

81. Wickham, H. ggplot2: Elegant Graphics for Data Analysis. (Springer International Publishing, 2016).

82. Zhu, A., Ibrahim, J. G. & Love, M. I. Heavy-tailed prior distributions for sequence count data: removing the noise and preserving large differences. Bioinformatics 35, 2084–2092, doi:10.1093/bioinformatics/bty895 (2019).

83. Quinlan, A. R. & Hall, I. M. BEDTools: a flexible suite of utilities for comparing genomic features. Bioinformatics 26, 841–842, doi:10.1093/bioinformatics/btq033 (2010).

84. Love, M. I., Huber, W. & Anders, S. Moderated estimation of fold change and dispersion for RNA-seq data with DESeq2. Genome Biol 15, 550, doi:10.1186/s13059-014-0550-8 (2014).

85. Edgar, R., Domrachev, M. & Lash, A. E. Gene Expression Omnibus: NCBI gene expression and hybridization array data repository. Nucleic Acids Res 30, 207–210, doi:10.1093/nar/30.1.207 (2002).

