## Supplementary Figures S1-S4 for "Sequential deregulation of histone marks, chromatin accessibility and gene expression in response to PROTAC-induced degradation of ASH2L"

[1]Institute of Biochemistry and Molecular Biology, Faculty of Medicine, RWTH Aachen University, Pauwelsstrasse 30, 52074 Aachen, Germany
[2]Contributed equally
[3]Present address: Bayer AG, Crop Science Division, R&D, Pest Control, 40789 Monheim am Rhein, Germany
[4]Present address: Institute of Human Genetics, Faculty of Medicine, University of Bonn, Venusberg-Campus 1, 53127 Bonn
[5]Retired

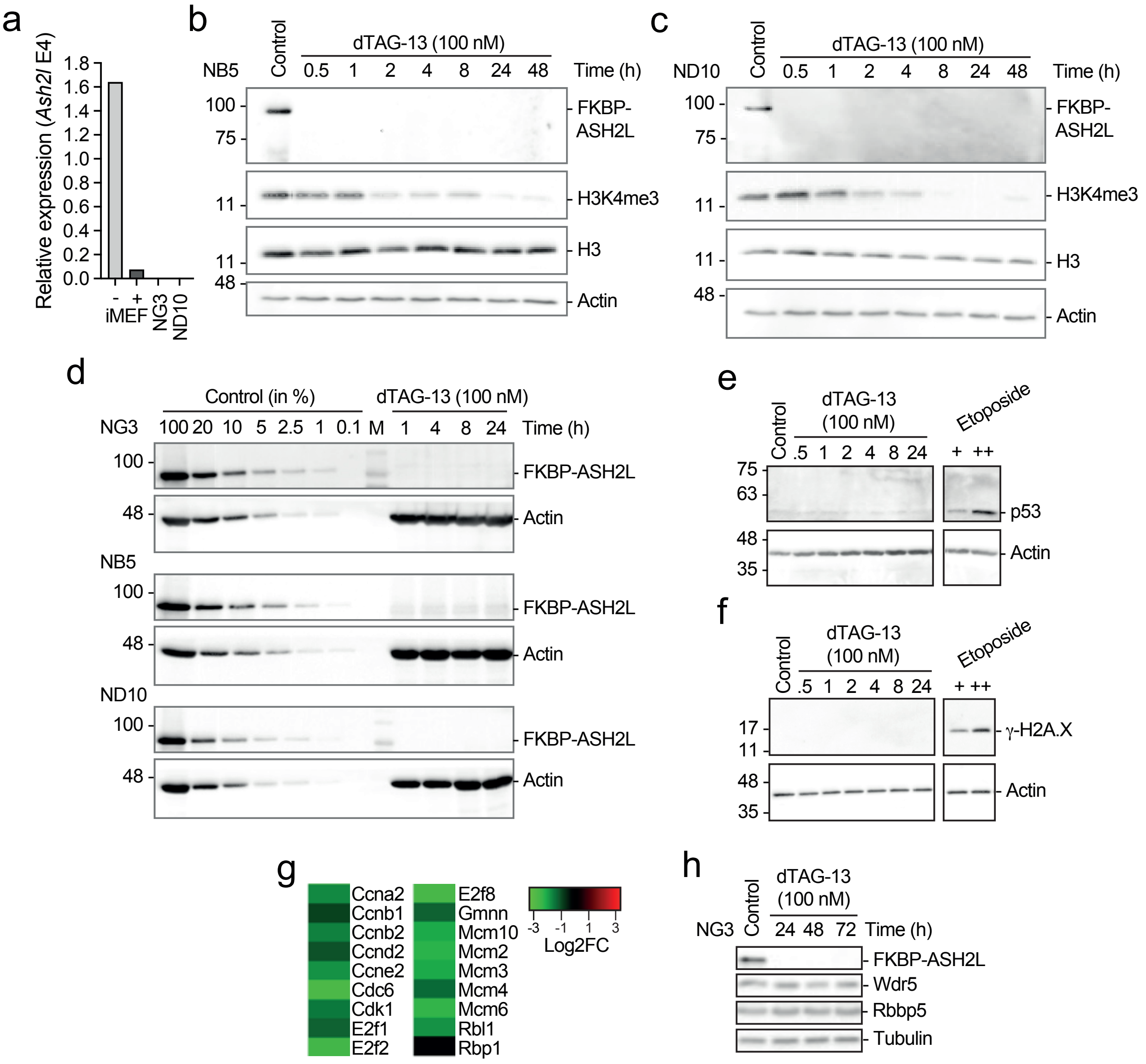


Supplementary Figure S1 (supporting Figures 1 and 2)

(a) RT-qPCR analysis of *Ash2l* transcripts from iMEF cells treated ±hydroxytamoxifen (-/+) or from NG3 and ND10 cell clones.

(b-d) FKBP-HA$_2$-ASH2L expressing cells were treated with dTAG-13. The indicated proteins were identified on Western blots, the cell clones analyzed are indicated.

(e and f) NG3 cells were treated with dTAG-13 as indicated. For control, cells were treated with etoposide (100 and 400 µM) for 4 hrs. p53 and γ-H2AX were stained as indicated.

(g) Down-regulated genes, which encode cell cycle regulators and replication factors, after dTAG-13 treatment for 24 hrs. (from Supplementary Table S1).

(h) Analysis of the WRAD components Wdr5 and Rbbp5 upon loss of FKBP-HA$_2$-ASH2L. Tubulin is shown for control.

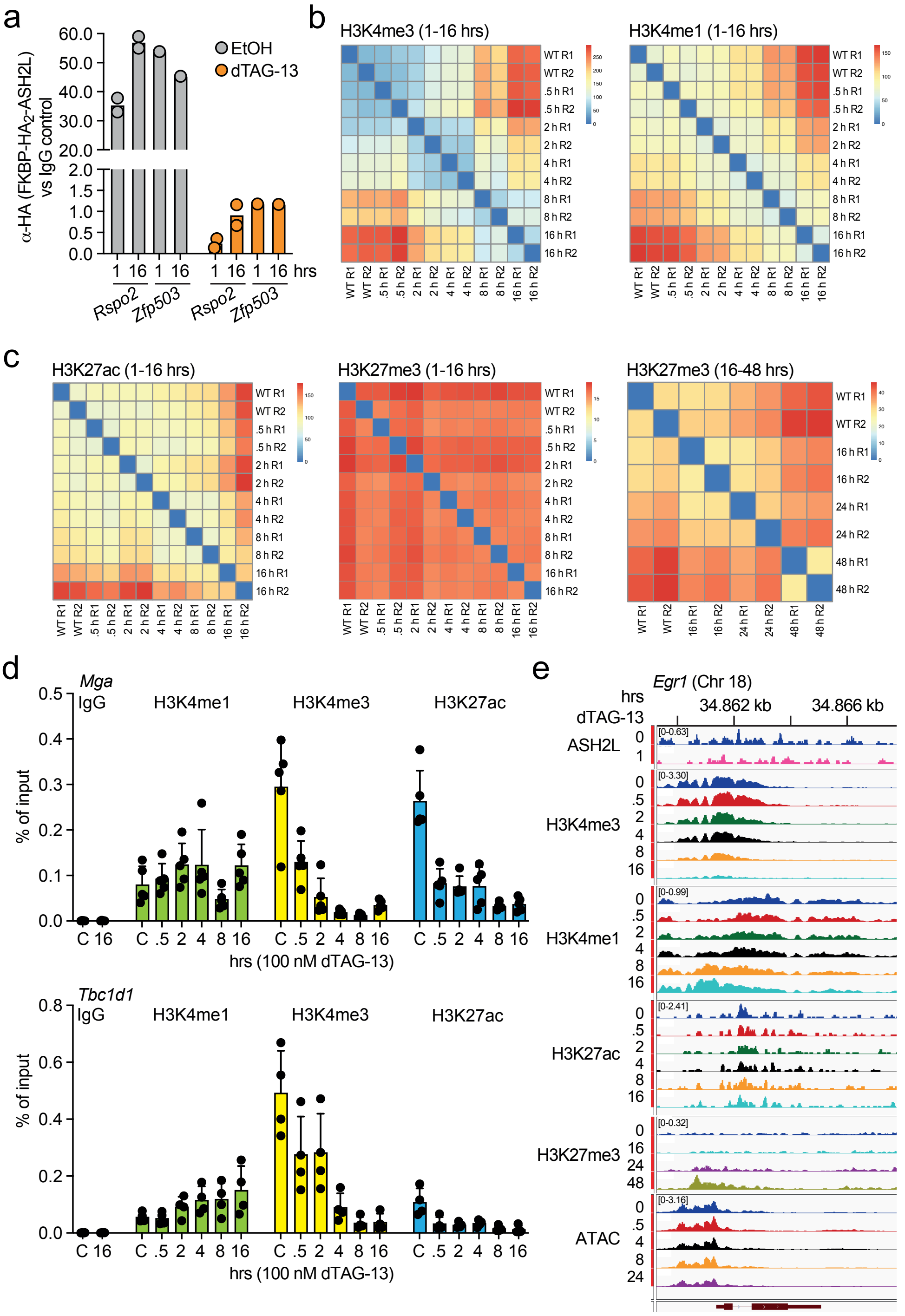


Supplementary Figure S2 (supporting Figure 3)

(a) Cells were treated with or without dTAG-13 (100 nM) for 1 or 16 hrs. ChIP-qPCR were performed using either HA-specific antibodies to immunoprecipitate FKBP-HA$_2$-ASH2L or control antibodies. Mean values of two replicates are displayed.

(b and c) Heatmaps showing sample-to-sample distances comparing all ChIP-seq samples of the indicated histone marks for both biological replicates to each other.

(d) ChIP-qPCR analyses of selected promoters for the indicated histone marks. Indicated are mean values ±SD of 4 to 5 measurements.

(e) Screen shots of integrative genomics viewer (IGV) of the *Egr1* promoter region showing the normalized BigWig tracks (normalized using counts per Million) of ChIP-seq experiments of the indicated histone marks and of ASH2L. Additionally, the chromatin accessibility is summarized as measured by ATAC-seq at the indicated timepoints after dTAG-13 treatment.

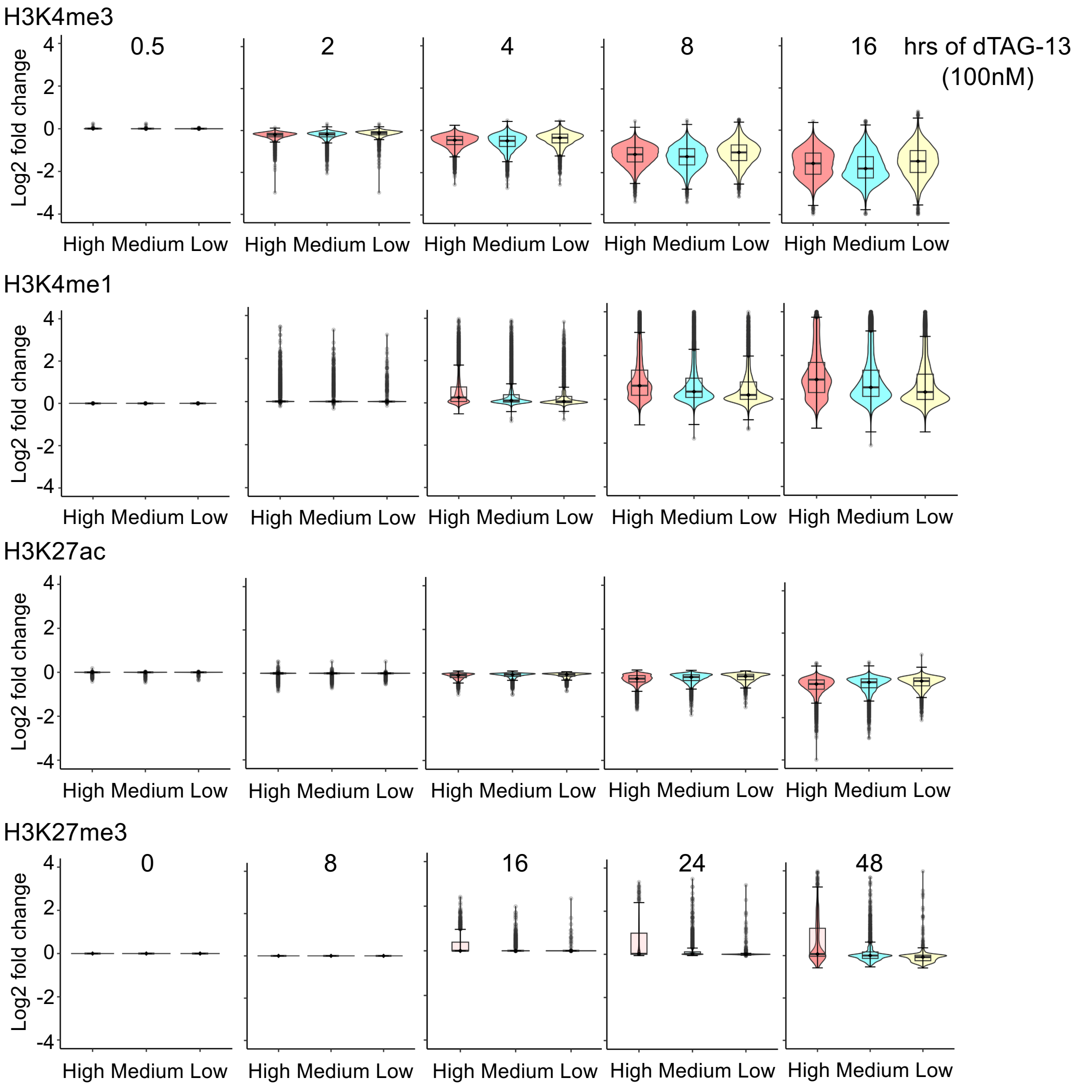


Supplementary Figure S3 (supporting Figure 4)
H3K4me3 binding sites (25431) were divided into three equal groups (8477 each). A window of ±1 kb around transcriptional start sites of all promoters was considered. The total number of binding sites within this window at promoters in the categories high, medium and low is 7937, 5413 and 1955, respectively. The log2 fold changes of signals of the indicated histone marks at the promoters of these three classes are displayed.

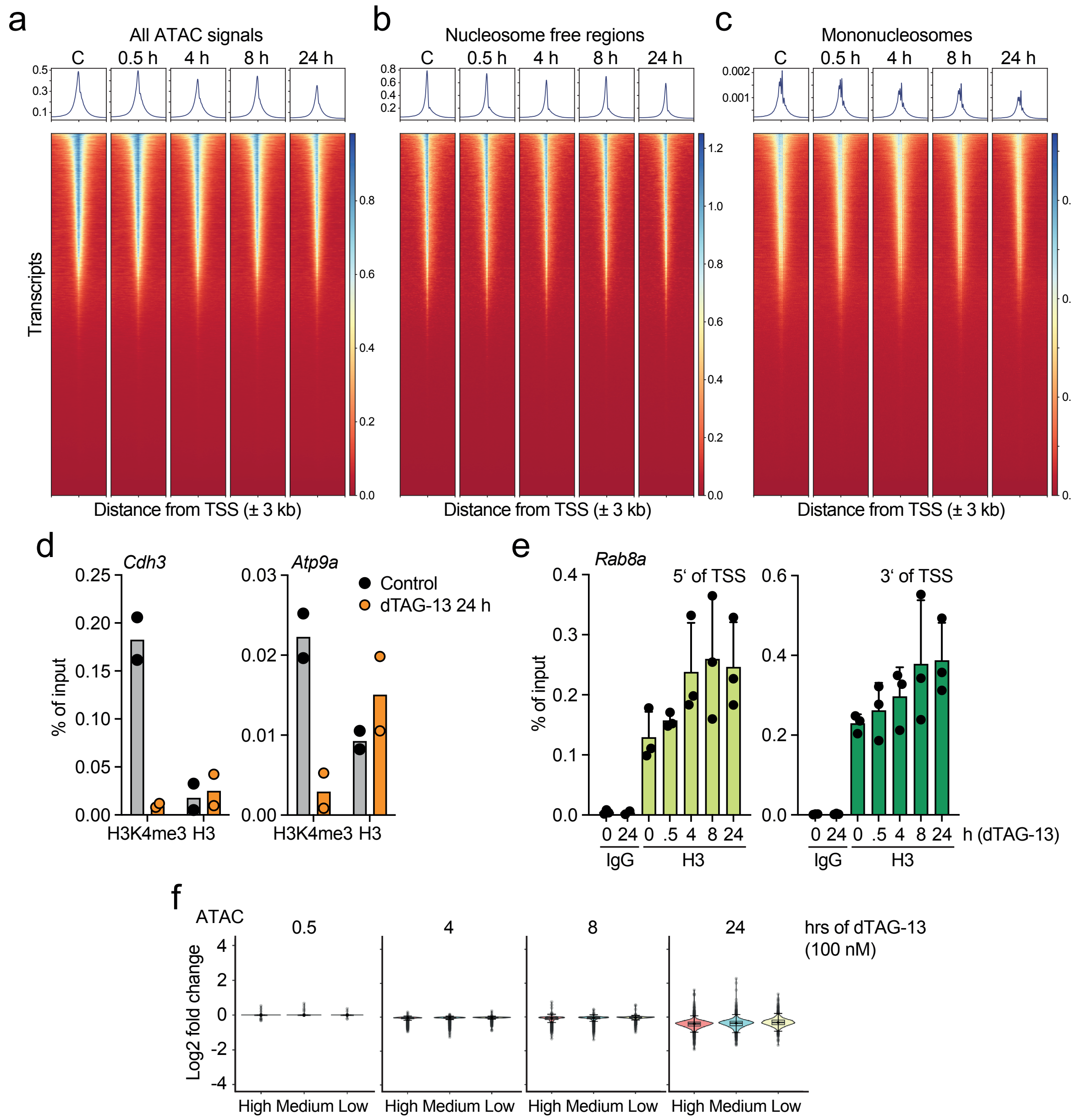


Supplementary Figure S4 (supporting Figure 6)

(a-c) Heatmaps and plot profiles generated using DeepTools showing the normalized ATAC-seq signals in response to dTAG-13 treatment at ±1 kb of transcriptional start sites (TSSs) of all annotated transcripts in mm10 (normalized using counts per Million). Nucleosome-free and mono-nucleosomes refer to ATAC-seq fragments that are smaller than 120 bp and between 130 and 200 bp, respectively.

(d and e) ChIP-qPCR measurements of H3K4me3 and histone H3. IgG was used for control. Between 2 and 3 experiments are displayed with mean values ±SD.

(f) Three equal groups of H3K4me3 signals were evaluated for being promoter associated (±1 kb of TSSs). This resulted in 7937, 5413 and 1955 sites in the three groups high, medium and low, respectively. The log2 fold changes of ATAC-seq signals at the promoters of these three groups are displayed.

**Supplementary Tables**

All sequencing data are available in NCBI's Gene Expression Omnibus as SuperSeries under accession number GSE241174.

**Table S1** List of significantly changed genes after 24h compared to the WT (RNA-seq)

**Table S2a** All identified Ash2l peaks across all the samples

**Table S2b** List of significantly changed Ash2l binding sites after 1h compared to the WT (ChIP-seq Ash2l)

**Table S3a** All identified H3K4me3 peaks across all the samples

**Table S3b** List of significantly changed H3K4me3 binding sites in each timepoint compared to the WT (ChIP-seq H3K4me3)

**Table S4a** All identified H3K4me1 peaks across all the samples

**Table S4b** List of significantly changed H3K4me1 binding sites in each timepoint compared to the WT (ChIP-seq H3K4me1)

**Table S5a** All identified H3K27ac peaks across all the samples

**Table S5b** List of significantly changed H3K27ac binding sites in each timepoint compared to the WT (ChIP-seq H3K27ac)

**Table S6a** All identified H3K27me3 peaks across all the samples (ChIP-seq H3K27me3_First experiment)

**Table S6b** List of significantly changed H3K27me3 binding sites in each timepoint compared to the WT (ChIP-seq H3K27me3_First experiment)

**Table S7a** All identified H3K27me3 peaks across all the samples (ChIP-seq H3K27me3_Second experiment)

**Table S7b** List of significantly changed H3K27me3 binding sites in each timepoint compared to the WT (ChIP-seq H3K27me3_Second experiment)

**Table S8a** All identified peaks across all the samples (ATAC-seq)

**Table S8b** List of sites significantly changed in accessibility in each timepoint compared to the WT (ATAC-seq)

**Table S9** List of all identified putative enhancers

**Table S10** List of significantly changed TF (ATAC-seq; differential analysis using RGT-Hint)

**Table S11** List of significantly changed genes after 4h or 8h compared to the WT (Nascent RNA; Click-it)
